# Post-training sleep modulates motor adaptation and task-related beta oscillations

**DOI:** 10.1101/2023.05.14.540662

**Authors:** Mohamed S. Ameen, Marit Petzka, Philippe Peigneux, Kerstin Hoedlmoser

## Abstract

Motor adaptation reflects the ability of the brain’s sensorimotor system to flexibly deal with environmental changes to generate effective motor behaviour. Whether sleep contributes to the consolidation of motor adaptation remains controversial. In this study, we investigated the impact of sleep on motor adaptation and its neurophysiological correlates in a novel motor adaptation task that leverages a highly automatized motor skill, i.e., typing. We hypothesized that sleep-associated memory consolidation would benefit motor adaptation and induce modulations in task-related beta band (13-30Hz) activity during adaptation. Healthy young male experts in typing on the regular computer keyboard were trained to type on a vertically mirrored keyboard while brain activity was recorded using electroencephalography (EEG). Typing performance was assessed either after a full night of sleep with polysomnography or a similar period of daytime wakefulness. Results showed improved motor adaptation performance after nocturnal sleep but not after daytime wakefulness, and decreased beta power (a) during mirrored typing as compared to regular typing, and (b) in the post-sleep vs. the pre-sleep mirrored typing sessions. Furthermore, the slope of the EEG signal, a measure of aperiodic brain activity, decreased during mirrored as compared to regular typing. Changes in the EEG spectral slope from pre- to post-sleep mirrored typing sessions were correlated with changes in task performance. Finally, increased fast sleep spindle density (13-15Hz) during the night following motor adaptation training was predictive of successful motor adaptation. These findings suggest that post-training sleep modulates neural activity mechanisms supporting adaptive motor functions.

## 1. Introduction

Motor adaptation refers to the process of adjusting automated motor skills based on sensory error signals in response to changes in environmental demands (Bastian, 2008). Not only is motor adaptation fundamental for the acquisition of new motor skills, but it is also important for the fine-tuning of existing ones through updating internal mental models of movements (Kawato, 1999).

The neural mechanisms underlying motor adaptation have been investigated, with recent Animal as well as human findings highlighting the role of oscillatory brain activity, specifically beta oscillations (12-30 Hz), in adjusting motor-related processes (Darch et al., 2020; Jahani et al., 2020). Indeed, beta band activity has been linked to several aspects of sensorimotor processing in the cortex, including motor planning, execution, and sensorimotor integration (Spitzer and Haegens, 2017). Decreased beta band power before movements reflects increased adaptive drive (Darch et al., 2020; Espenhahn et al., 2019). However, the mechanisms governing the consolidation of these adaptive changes, particularly within the context of sleep, remain poorly understood.

In recent years, a growing body of evidence has highlighted the unique contribution of sleep to motor adaptation consolidation (Bothe et al., 2019, 2020; Hoedlmoser et al., 2015; Huber et al., 2004; Plihal & Born, 1997). Indeed, previous studies suggested that the increase in the density of fast sleep spindles (13-15 Hz), observed during stage 2 Non-rapid eye movement (NREM) sleep after training support motor adaptation (Bothe et al., 2020; Solano et al., 2022). Furthermore, rapid eye movement (REM) sleep has also been associated with the consolidation of motor adaptation (Bothe et al., 2018; for an overview see King et al., 2017). Interestingly, studies have shown an increase in beta power during REM sleep in the Anterior Cingulate Cortex, a crucial brain region involved in the adaptation of motor output (Ebitz & Hayden, 2016; Vijayan et al., 2017). Several other studies, however, found no effect of sleep on motor adaptation, suggesting that time, whether spent awake or asleep, is the limiting factor for the consolidation of motor adaptation (Debas et al., 2010; Doyon et al., 2009; Thürer et al., 2018). As a result, the specific influence of sleep on motor adaptation as well as its role in modulating beta oscillations in that context have not been established.

In this framework, the present study aimed at investigating whether (1) sleep benefits the adaptation of motor skills and (2) does so through modulating power beta oscillations during the motor adaptation task. To this end, we designed a novel motor adaptation task leveraging an already automatized motor skill, i.e., typing. Specifically, we trained human experts in touch-typing on a computer keyboard to type on a vertically mirrored keyboard, therefore requiring a consistent motor adaptation, and measured their mirrored typing performance before and after a retention interval of either nocturnal sleep (∼8h with polysomnography; PSG), or a similar period of daytime wakefulness.

We hypothesized to observe a beneficial role of post-learning sleep in motor adaptation that is reflected in decreased task-related beta power following sleep. We anticipated a correlation between the decreased task-related beta power from the pre-sleep to the post-sleep adaptation and increased beta power during REM sleep as well as increased fast spindles density. Using multivariate decoding techniques, we posited that sleep-associated motor adaptation consolidation would result in increased similarity of brain activity between the original task and the adaptation task following sleep, evidencing successful sleep-associated memory consolidation. Finally, to probe the cortical processes occurring during motor adaptation, we explored the relationship between the spectral slope of the EEG signal and motor adaptation. The EEG slope is a measure of the balance between cortical excitation and inhibition (Gao et al., 2017; Lombardi et al., 2017) and may provide valuable insights on the excitation-inhibition ratio during motor adaptation.

## 2. Methods

### 2.1. Participants

33 healthy male subjects (age range 19-29 years, mean = 25 ± 4 years) were recruited for this study that was approved by the ethics committee of the University of Salzburg. The decision to include only male participants was initially taken to minimize variations in motor performance and avoid any task-related effects due to differences in the menstrual cycle phase (Plamberger et al., 2021). The participants were randomly divided into a sleep group (n=16) and a wake group (n = 17) in a randomised, between-subjects design. We recruited experts in touch typing on the keyboard (250 correct keypresses/minute). Their expertise was estimated by measuring the time and accuracy of typing a specific text on the regular computer keyboard. The decision of the typing criterion assumes that experts in touch-typing are able to type at least 50 five-letter words per minute (Inhoff, 1991). Selection criteria were being male, right-handed, German-speaking, neural chronotypes, i.e., no extreme morning or evening types, with no history of severe organic and/or mental illness, no sleep disorders (i.e., Pittsburgh Sleep Quality Index Global Score > 5, Buysse et al., 1989), no signs of mood disorders (i.e., Self-Rated Anxiety Scale Raw Score < 36 and Self-Rated Depression Scale Raw Score < 40, no unusual bedtimes and/or extreme chronotype. Students received ECTS (European Credit Transfer System) points or monetary compensations for participating. They were informed about the aspects of the project in detail before they gave their written informed consent before the beginning of the study.

### 2.2. Experimental design (Fig. 1)

#### 2.1.1. Protocol (**Fig. 1A**)

Sleep-wake rhythm was monitored via wrist actigraphy starting three days prior to the first laboratory visit and for the whole duration of the experiment. Subjective sleep/wake rhythm and sleep quality was also collected via sleep logs (Saletu et al., 1987). After the entrance examination (day 1), participants in the sleep group slept for a baseline night on day 4 in the laboratory. The training session took place either in the evening (08:00pm - 11:00pm) of the 6th day for the sleep group, or in the morning (09:00am - 12:00pm) of the 4th day for the wake group. A retention period that included either 8h of sleep during the night at habitual bedtime (11:00pm to 07:00am) for the sleep group (testing starts at 09:00am) or a period of daytime wakefulness (approximately 8h) for the wake group (testing at 05:00pm to 07:00pm), preceded the testing session Vigilance was monitored with a 10 minute visual psychomotor vigilance task (PVT; adapted from Dinges & Powell, 1985) before and after training as well as after the retention interval. Participants in the wake group were instructed to a) not sleep (controlled via wrist actigraphy), and b) not type on any computer keyboard during the retention interval.

**Figure 1.**
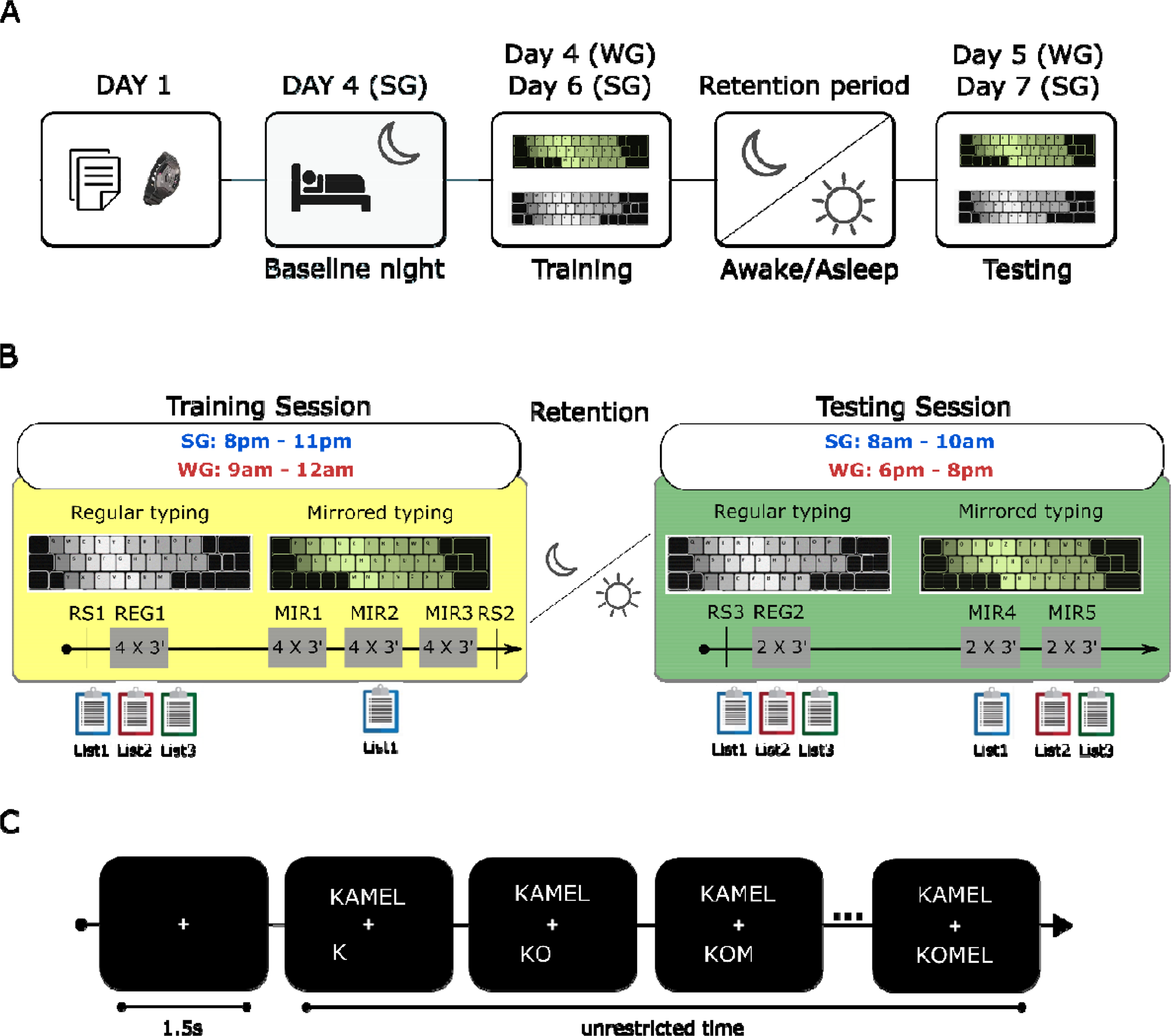
Experimental design. A) Experimental protocol: after the entrance examination, selected participants were divided into a sleep (SG) and a wake (WG) group. In the evening of day 4 for the sleep group, participants spent a baseline night with polysomnography (PSG) in the laboratory for habituation. The experimental procedure started with a training session followed by a retention interval that included either eight hours of nocturnal sleep or a similar period of daytime wakefulness. After the retention interval, participants performed the testing session. B) Paradigm: (Left) The training session started with a resting state recording (RS1) followed by four typing blocks on the regular keyboard (REG1). After a 15-minute break the mirrored typing session started with three mirrored typing sessions (MIR1, MIR2, and MIR3) which consisted of four three-minute typing blocks. (Right) The testing session started with regular typing (REG2) which consists of two typing blocks of three minutes. After a 5-minute break, participants performed two mirrored typing blocks (MIR4 & MIR5). IN REG1 and REG2 participants typed all the 18 words, in MIR3 and MIR4, participants typed the six words in List1, while in MIR5 they typed the rest of the 18 words (12 words; List2 and List3). C) Typing trial structure. The trials started with a fixation cross for 1.5 seconds, then a word appeared on the screen above the fixation cross. The word remained on the screen until the participant typed five letter regardless of whether they were correct. Participants received immediate feedback by seeing the letters they typed below the fixation cross. However, there was no option to correct what has been typed. There were no time constraints for typing, as the trials ended when participants typed five letters regardless of whether they were correct. REG1: Regular typing before retention, + p < 0.1, * p < 0.05. REG2: Regular typing after retention, MIR1-3: Mirrored typing before retention, MIR4-5: Mirrored typing after retention.

#### 2.2.1. Procedure (**Fig. 1B**)

Participants had to type five-letter, German words on a regular QWERTZ computer keyboard (REG) as well as on a vertically mirrored keyboard (MIR) as rapidly and as accurately as possible. The mirrored keyboard was the regular QWRETZ keyboard with the output of the keys vertically mirrored (see Fig. S1A). The 18 German words were composed of 17 different letters (A, C, D, E, F, H, I, K, L, M, N, O, P, R, S, T, U) and had to be typed bimanually: two-to-three letters with the left/right hand. 10-minutes resting state EEG recording (RS1; 5 minutes recording with eyes open then 5 minutes with eyes closed). Thereafter, the first regular typing (REG1) session started. REG1 consisted of four three-minute blocks of typing on a regular keyboard, separated by 15s breaks. During these typing-blocks (4×3min), participants had to type all of the 18 words. After a ten-minute break, they were trained on typing a subsample of the words (*List 1;* six words composed of eight different letters: A, E, K, L, M, N, P, R) on the vertically mirrored keyboard. On this mirrored keyboard, participants had to learn to perform the same sequence of key presses, but with the other hand (see Fig. 1B and Fig. S1). The mirrored typing learning consisted of three sessions (MIR1-MIR3). Each session was composed of four three-minute blocks, separated by 15s breaks, summing up to a total of twelve three-minute blocks of training on the mirrored keyboard. During MIR1-MIR2, the structure of the mirrored keyboard was displayed at the bottom of the screen to aid the learning process. For MIR3, however, participant had to type without any visual aid. At the end of the training session, another resting state (RS2) recording took place. Following motor adaptation on the mirrored keyboard, participants spent a retention interval that included either 8h of wakefulness during the day or 8h overnight sleep. After the retention interval, the testing session started with a final resting state recording (RS3), followed by two three-minute blocks of regular typing (REG2 – 2×3min; 18 words), then two three-minute mirrored typing blocks (MIR4-MIR5 2×3min). During MIR4 participants had to type the six words that were used for the training. During MIR5, they had to type the 12 words that were not included in the mirrored keyboard training. Six of those 12 words that were presented during MIR5 consisted of the same letter as the trained words (*List 2*; A, E, K, L, M, N, P, R). The other six words were composed of nine different letters (*List 3;* C, D, F, H, I, O, S, T, U) which were not practiced on the mirrored keyboard in the training session before retention. For more information on the word lists, see Fig. S1B. **2.2.3. Trial structure (Fig. 1C):** Each trial started with a fixation cross (1.5s), after which a word appeared above the fixation cross. For the trial to end, participants had to press five keys regardless of whether they were correct. The time taken to type the five letters was computed starting from the first button press until the fifth. There was no time limit for the trials. Participants received immediate feedback as they saw the letter they typed on the screen; however, they had no option to change it. The task was designed in that way to measure both typing accuracy and speed.

### 2.3. Behavioural analysis

We derived a behavioural estimate of typing performance accounting for both typing accuracy and typing speed by dividing typing-accuracy (ratio of correct letters per five-letter word) by typing-speed (time in seconds required to type the five letters of the word). Typing performance was measured separately for word in the session.

### 2.4 EEG acquisition

EEG during typing and as well as during sleep was recorded using a 32-channel Neuroscan system (Scan 4.3.3 Software, Neuroscan Inc.; Charlotte, USA) with 500Hz sampling rate and channel Cz as online reference. We used a montage with 11 EEG electrodes (F3, F4, Fz, C3, C4, Cz, P3, P4, Pz, O1, and O2). Eye movements were recorded via one vertical and one horizontal electrooculogram (EOG) channel, and muscle activity via a bipolar submental electromyogram (EMG) channel. Ground electrode was placed at AFz location. Impedances were kept below 5kΩ.

### 2.5. Sleep staging and spindle detection

Sleep was automatically scored on 30s epochs using the Siesta group (Somnolyzer 24×7; The SIESTA Group Schlafanalyse GmbH., Vienna, Austria; Anderer et al., 2005; Anderer et al., 2010). Afterwards, sleep was visually inspected by an expert scorer according to the criteria of the American Academy of Sleep Medicine (AASM, American Academy of Sleep Medicine & Iber, 2007). Similarly, sleep spindles were detected using the automatic SIESTA algorithm (ASK analyser, The Siesta Group Schlafanalyse GmbH., Vienna, Austria, Gruber et al., 2015). First, all electrodes were referenced to the average of both mastoids before filtering the signal between 0.1 and 40 Hz and resampling it to 128Hz. The algorithm first filters the raw data between 11 and 16 Hz before detecting spindle events using the criteria described in Schimicek et al. (1994). Only events with an amplitude >12µv and duration between 500ms and 3000ms were considered. The detected spindles were compared to a template generated based on the visual scoring of experts on 8730 minutes of PSG data from 189 healthy participants and 90 participants with sleep disorders via Linear discriminant analysis (LDA). Only events with an LDA score of 1.7 or higher corresponding to 98% specificity were considered for the analysis (for more details see: Anderer et al., 2005). Only those spindles that occurred during N2 and N3 sleep were included. The frequency of each sleep spindle was measured using period-amplitude analysis of the band-pass filtered signal in the time domain and spindles were subdivided into slow (11-13 Hz) and fast (13-15Hz) spindles. Spindle density was calculated as the number of events per minute of N2, N3, or both (NREM).

### 2.6. EEG data pre-processing and analysis

EEG data were pre-processed using the EEGLab toolbox (v14.1.1b) implemented in Matlab (v2019a). Raw data were notch-filtered at 50Hz to remove line noise using the CleanLine toolbox in EEGLab, high-pass-filtered at 0.1 Hz, and then referenced to an average reference. For our analysis, we divided each session into 10s epochs with five seconds of overlap. The epoched data were automatically inspected for artefacts and bad epochs with large amplitude fluctuations (above 1000µV) were removed. On average 6.32 ± 0.82 % of trials (7.52 ± 2.24 trials) per participant per condition were removed. Time-frequency analyses were performed in Fieldtrip (Oostenveld et al., 2011). Following pre-processing, the spectral power of the epoched data was estimated using a multitaper fast Fourier transform (FFT) with discrete prolate spheroidal sequences (DPSS) tapers. We used frequency smoothing of 0.4Hz from 0.5Hz to 45Hz in 0.5Hz steps and 100ms temporal steps fitting 5 cycles per time window. We used large epochs of 10s to avoid any contamination due to edge artifacts. For keypress-related analysis, the epochs were centred on the keypresses. We separated periodic and aperiodic components (except for MVPA analysis) of the EEG signal using the irregular resampling auto-spectral analysis (IRASA, Wen & Liu 2016) implementation in Fieldtrip. The spectral slope of the EEG signal was estimated by calculating the slope of the fractal component of the signal in the log-log (log frequency, log power) space. We chose a broadband range for slope estimations, i.e., 1-45Hz, as a compromise to achieve the best possible representation of the signal without including any noise in the estimation due to line noise or line-noise-removal procedure.

### 2.7. Multivariate pattern analysis (MVPA)

To highlight EEG activity differences and commonalities between regular and mirrored typing, we performed multivariate decoding using the MVPA-Light toolbox (Treder, 2020). First, an LDA classifier was trained on a subset of the data to distinguish between the conditions. The pre-processing and time-frequency estimation procedure are the same as described in section 2.7. In addition, the pre-processed EEG signal was z-scored, and underwent K-fold cross-validation (K=five, two repetitions). We performed (spectral) decoding on the central electrodes (C3 and C4) and on 600ms epochs centred on keypresses. For spectral decoding, we used time-frequency transformations (section 2.6) and tested the ability of the classifier to decode between the conditions based on each single frequency (0.5-45Hz in 0.5Hz steps) at each time point. For such analyses, we did not perform IRASA-based 1/f correction for this analysis, as we wanted to maintain the time dimension which is discarded in the process of IRASA 1/f calculation. For the decoding results, we report the area under the curve (auc) parameter, as it is less sensitive than the decoding accuracy parameter to class imbalances thus mitigating the effect of the difference in the number between regular and mirrored typing trials.

### 2.8. Statistical analysis

Robust statistical analysis was performed using rank-based non-parametric tests as implemented in the *nparLD* package (Ngouchi et al., 2012) available in R (v4.1.2). It reports ANOVA-type statistics (ATS), p-values (alpha = 0.05, two-sided), as well as relative treatment effects (RTE) as effect sizes. RTE values represent the probability of the values from the whole dataset being smaller than a randomly chosen observation from the respective group. The higher the RTE value of one condition means a higher probability that a randomly chosen value from that condition is larger than that randomly drawn from the whole dataset, and vice versa. We performed post-hoc tests via the ‘*NparLD*’ function with Bonferroni’s correction for multiple comparisons. For all time-series analyses, non-parametric permutation statistics were performed as implemented in Fieldtrip (Maris and Oostenveld, 2007). Two-sided (un)paired-sample t-tests were computed with a Monte Carlo procedure over 5000 permutations (cluster-alpha = 0.05 and critical alpha = 0.025). The sum of the t-values (∑t) is reported, as well as Cohen’s d effect sizes calculated over all possible permutations. For correlations, we used Pearson’s correlation in case of normally distributed data and Spearman as indicated by the Shapiro-Wilk test. For the topographical plots, Spearman’s method was applied whenever the normality assumption was violated in at least 1 electrode. Rho and p-values are reported; in case of multiple correlations, correction was done using Monte-Carlo procedure with 10000 permutations as implemented in fieldtrip. These analyses were pre-registered and can be accessed at the following link: https://osf.io/jfdv6.

## 3. Results

### 3.1. Nocturnal sleep benefits the adaptation of fine motor-skills

A mixed 2×2 mixed non-parametric test on mirrored typing performance (Fig. 2A) with a within-subject factor *Time* (before vs. after retention), and a between-subjects factor *Group* (Sleep vs. Wake) revealed a main effect of *Time (ATS(1) = 22.61, p < 0.001, RTE_pre_ = 0.45, RTE_post_ = 0.55*) with higher mirrored typing performance after than before the retention interval for both groups. The main Group effect was non-significant (*ATS(1) = 1.71, p = 0.19, RTE_Wake_ = 0.44, RTE_Sleep_ = 0.57*). A significant *Group x Time* interaction effect (*ATS(1) = 7.04, p = 0.007, RTE_preWake_ = 0.42, RTE_postWake_ = 0.46, RTE_preSL_ = 0.49, RTE_post_ = 0.64*) showed that performance significantly improved after nocturnal sleep (*ATS(1) = 25.27, p_bonf_ < 0.001, RTE_pre_ = 0.42, RTE_post_ = 0.58*) but did not change following daytime wakefulness (*ATS(1) = 1.63, p_bonf_ = 0.4, RTE_pre_ = 0.48, RTE_post_ = 0.52*).

**Figure 2.**
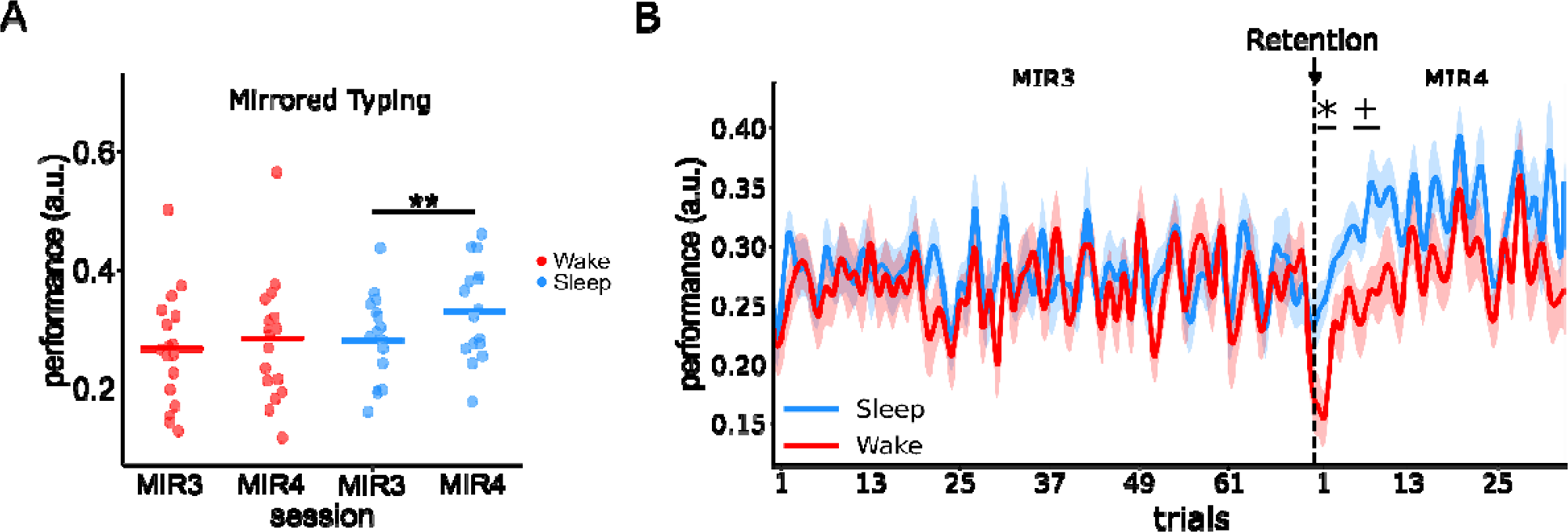
Behavioural results. A) Typing performance on the mirrored keyboard before and after the retention interval averaged over each session. Typing performance on the mirrored keyboard significantly improved only after sleep (MIR3 to MIR4). There was no significant change after wakefulness (see also Fig. S2). B) Trial-by-trial (word-by-word) analysis of typing performance on the mirrored keyboard showing that the sleep group performed significantly better than the wake group at the early phase of the post-retention session (first eight trials). REG1: Regular typing before retention, + p < 0.1, * p < 0.05. REG2: Regular typing after retention, MIR3: Mirrored typing before retention, MIR4: Mirrored typing after retention.

When the regular-typing trials were included in the statistical model, the results confirmed that sleep-dependent positive effects were specific to mirrored typing with no interference with regular typing performance and no effect of wakefulness on performance in both typing conditions (Fig. S2A).

To assess whether the effect of post-training sleep on motor adaptation was transient or rather sustained throughout the whole testing session, we compared performance on the mirrored keyboard between the groups (sleep vs. wake) on a word-per-word basis (Fig. 2B). Results revealed that the highest between-group difference in motor adaptation performance in the first eight trials (out of approximately 30 trials) after the retention interval (trials 1-2: ∑*t(31) = 6.91*, *p = 0.03*, *d= 0.56*; trials 6-8: ∑*t(31)* = 5.49, *p = 0.023*, *d = 0.94*). Noticeably, performance decline in the wake group was visible immediately after the retention interval, an effect that was absent in the sleep group. We found no between-group differences in regular typing performance after sleep (see Fig. S2B-D).

### 3.2. Central beta power during adaptation decreased following nocturnal sleep

To characterize the changes in central beta power during typing from before to after the retention interval, we compared beta power during the typing sessions before (REG1 and MIR3) and after the retention interval (REG2 and MIR4). We performed a 2×2x2 mixed non-parametric test with within-subject factors *Keyboard,* and *Time*, and a between-subjects factor *Group* (Fig. 3A and see Table 1 for statistical results). Central beta power during mirrored typing was marginally lower than that during regular typing. However, decreased beta power from regular to mirrored typing becomes significant when looking at the global power averaged over all EEG electrodes (Fig. S3A). There was a significant effect of *Time* as beta power decreased after the retention interval for both groups as compared to before the retention interval, and a significant effect of *Group* as beta power was lower in the sleep than the wake group. Moreover, there was a significant interaction *Group x Time* interaction and post-hoc tests revealed that central beta power decreased after nocturnal sleep (*ATS(1) = 31.38, p_bonf_ < 0.001, RTE_pre_ = 0.56, RTE_post_ = 0.44*) but did not change following a period of daytime wakefulness (*ATS(1) = 1.13, p_bonf_ = 0.58, RTE_pre_ = 0.49, RTE_post_ = 0.51*). *Group x Keyboard*, *Keyboard x Time* and *Group x Time x Keyboard* interactions were not significant. Contrary to our hypothesis, however, the difference in central beta power from pre- to post-sleep did not correlate with the change in performance in the adaptation task (Fig. S3B). To make sure that differences in beta power were not due to circadian variations, we compared beta oscillatory power in the eyes-closed resting state recording before training (RS1) and before testing (RS3). Beta power increased from RS1 to RS3, i.e., in the opposite direction of the task-induced beta power changes ((*ATS(1) = 24.82, p < 0.001, RTE_SLpre_ = 0.35, RTE_SLpost_ = 0.54, RTE_Wpre_ = 0.42, RTE_Wpost_ = 0.68*, see supplementary material and Fig. S3C). For more information on the temporal aspects of beta power fluctuations in relation to keypresses please refer to Fig. S4.

**Figure 3.**
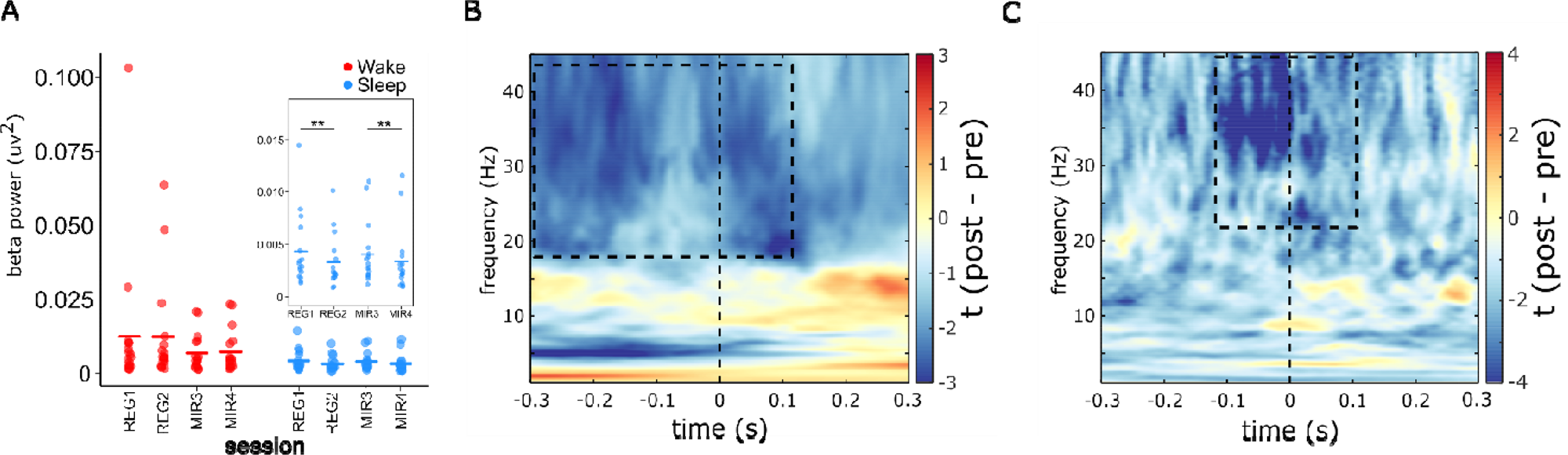
Sleep induced changes in adaptation-related brain activity. A) Beta band power averaged over electrodes C3 an C4 during regular and mirrored typing before and after retention. Central beta power decreased during mirrored typing. After sleep beta power decreased during both regular and mirrored typing but increased after a period of wakefulness. The box magnifies beta power during mirrored typing for a better illustration of power differences. B) T-map depicting the difference in the classification (regular typing vs. mirrored typing) from pre-sleep to post-sleep. The accuracy of the classifier to differentiate between regular and mirrored typing using beta band power decreased after sleep (-0.3s – 0.1s relative to keypresses). C) T-map depicting the difference in the classification accuracy of (correct vs incorrect) mirrored typing trials from before to after sleep (MIR4-MIR3). Note a decrease in decoding performance following sleep in the high beta range (>25Hz) and in the -0.1 -0.1s time window. REG1: Regular typing before retention, REG2: Regular typing after retention. MIR3: Mirrored typing before retention, MIR4: Mirrored typing after retention. **p < 0.001. Each point in panel A represent one subject and the crossbars represent the mean of the group. Vertical line at zero indicates button press. Dashed boxes indicate significant clusters. Dashed horizontal line in panel A represent chance level.

**Table 1.**
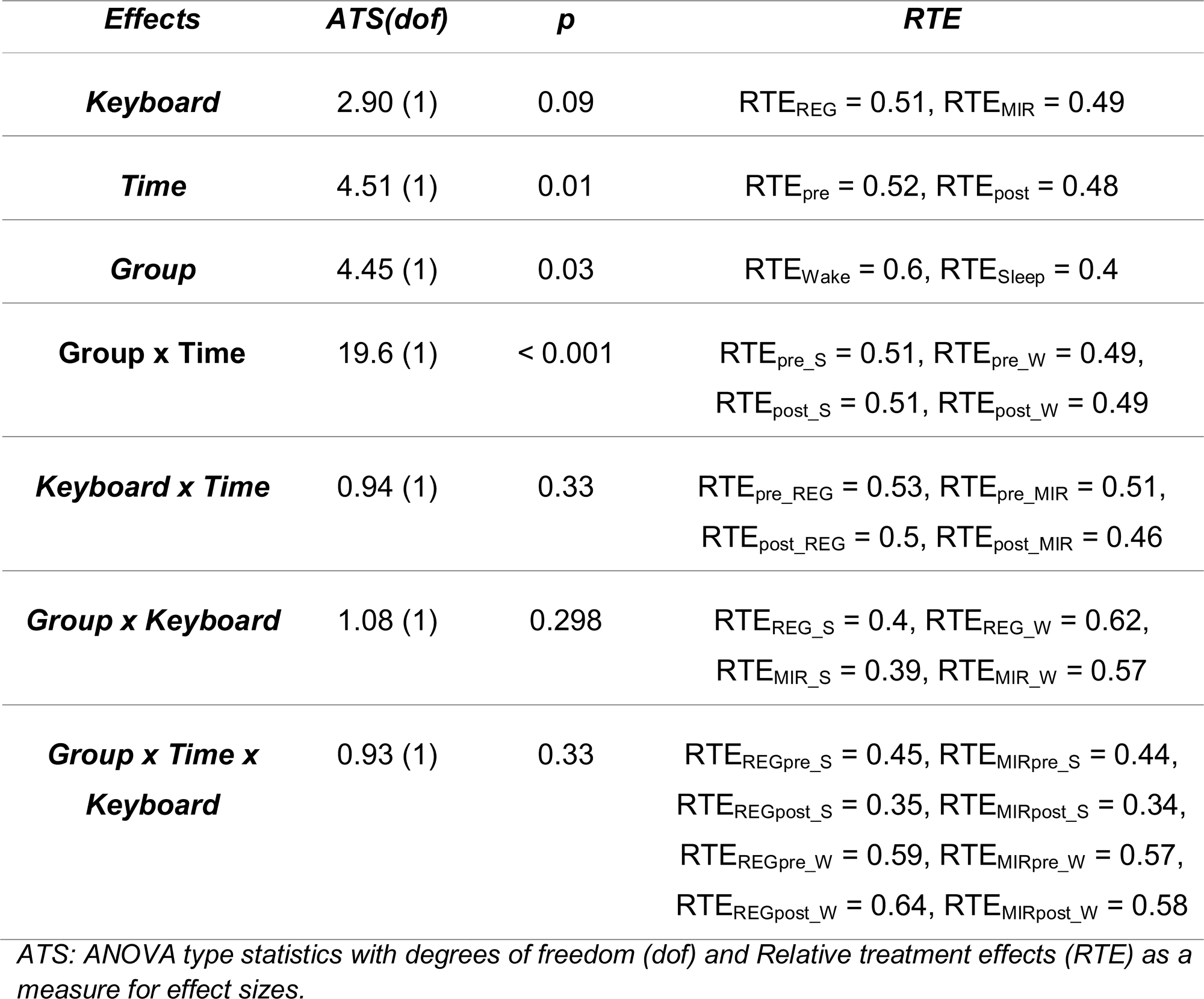
Statistical results for the difference in central beta power.

Next, we sought to investigate any differences in brain activity between mirrored and regular typing, and whether sleep modulated such differences. We hypothesized that sleep-associated consolidation would minimize differences in beta band activity resulting in neural patterns becoming more similar between mirrored and regular typing after sleep. To test this hypothesis, we employed multivariate decoding on the time-frequency data. That is, we trained a classifier on the spectral data of all typing trials (Regular and Mirrored) and tested its ability to differentiate (i.e., decode) between regular and mirrored typing trials based on spectral power. We then compared the decoding accuracy (i.e., the auc metric) before and after the retention interval (see section 2.7 for more details). Results show that the decoding accuracy in the beta range (> 18Hz) significantly dropped after sleep (Fig. 3B; -0.3s – 0.1s, ∑*t(15) = -5905.22, p = 0.008, d = -0.83*) but not after wakefulness (Fig. S5A), indicating that beta power became less differentiated between regular and mirrored typing only following sleep. Examination of the classifier results separately before and after sleep revealed that the highest accuracy (auc) in differentiating between regular and mirrored typing was observed in the beta frequency range (Fig. S5B). Consequently, we asked whether sleep time-frequency decoding would also be able to classify correct vs. incorrect mirrored typing trials. Therefore, we trained the classifier on the spectral data of all mirrored typing trials (correct and incorrect) and tested its ability to differentiate between correct and incorrect ones. Results revealed that the although the classifier was able to differentiate between correct and incorrect trials in the training session, this ability dropped in the post-sleep session (Fig. S5D). The highest decreasing in decoding was observed in the high beta (>25Hz) band after sleep around -1.1s – 0.9s relative to keypresses (Fig. 3C; ∑t(15) = -2803.67, p = 0.006, d = - 1.103). This effect was not observed after wakefulness (Fig. S5C).

### 3.3. NREM stage 2 fast sleep spindle density predicts motor adaptation

We then asked whether the change in fast spindle density from the baseline night to the experimental night correlates with the change in adaptive performance. Hence, we focused on central fast spindle densities (averaged over C3 and C4) in correspondence to our beta power analysis. There was a positive correlation between the change in fast spindle density during NREM stage 2 (N2) sleep from the baseline night to the experimental night, and performance gains on the mirrored keyboard from pre- to post-sleep (Fig 4A*; r(14) = 0.58, p = 0.03*). A cluster-based permutation analysis on Spearman correlations across electrodes (10000 permutations) highlighted a significant correlation over a cluster of 10 electrodes (Fig. 4B; all electrodes except O2; ∑*t(15) = 33.32, p < 0.001, d = 0.29*). We observed a similar pattern in N3 sleep and when combining N2 and N3 spindles (see Fig. S6A-D). Finally, there was no significant correlation between N2 fast spindle density and beta power change (*r(14) = -0.38, p = 0.14,* Fig. S6E). Furthermore, we were not able to confirm our hypothesis that REM density and REM beta power benefit motor adaptation (see Fig. S7A-B) as there was no change in REM density (time in REM / total sleep time) or beta band power during REM from the adaptation to the experimental night.

**Figure 4.**
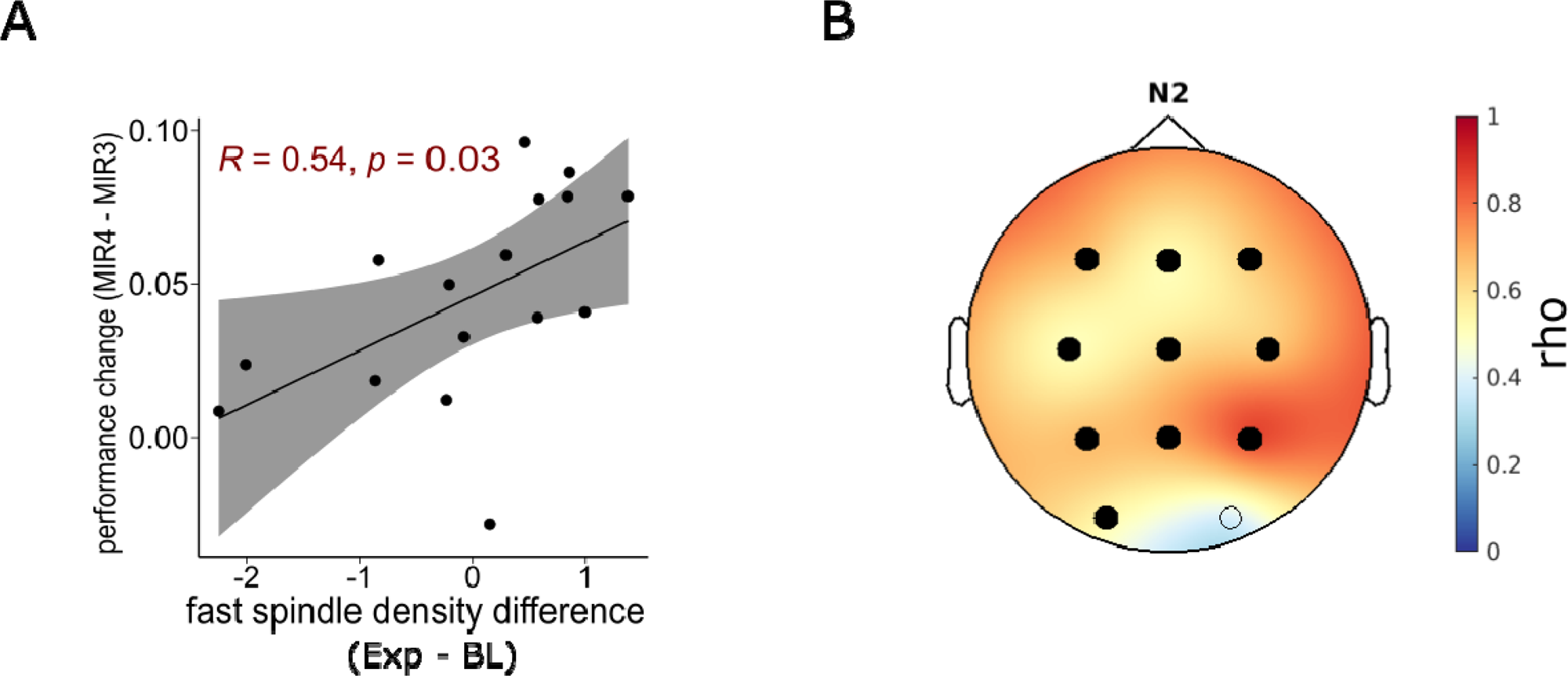
NREM stage 2 fast sleep spindle density predicts motor adaptation. A) Change in the density of central fast sleep spindles (averaged over C3 and C4) during NREM stage 2 sleep from the baseline to the experimental night correlated with performance change on the mirrored keyboard from pre- to post-sleep (MIR4 - MIR3). B) Correlation between fast spindle density and performance change significant over a cluster of 10 electrodes (all EEG electrodes except O2). BL: baseline night, Exp: experimental night. R: Pearson’s correlation. rho: Spearman’s correlation.

### 3.4. The spectral slope of EEG signals predicts motor adaptation

Finally, we performed an analysis to characterise changes in the spectral slope during motor adaptation. Typing sessions were segmented into 10s segments with five seconds of overlap. A 2×2x2 mixed non-parametric test with the slope of the 1-45Hz EEG signal averaged over all electrodes as the dependent variable and two within-subject factors *Keyboard* and *Time*, and a between-subjects factor *Group was performed (*see Table 2 for full statistical results). Results revealed a main *Keyboard* effect *(p = 0.009)* as the slope was steeper during mirrored than regular typing (Fig. 5A). There were no effects of *Time (p = 0.1)*, nor *Group (p = 0.68)*. There were also no significant interactions *Time x Group (p = 0.2)*, *Keyboard x Time (p = 0.2), Keyboard x Group (p = 0.3),* nor *Keyboard x Time x Group (p = 0.92)*. Additionally, we observed a significant positive correlation between the change in the spectral slope from the pre-sleep mirrored typing session (MIR3) to the post-sleep mirrored typing session (MIR4) and the change in task performance MIR4-MIR3 (Fig. 5B; *r(14) = 0.57, p = 0.02*). In the wake group, however, the changes in the spectral slope mediated by a period of wakefulness did not predict performance change (Fig. S7C).

**Figure 5.**
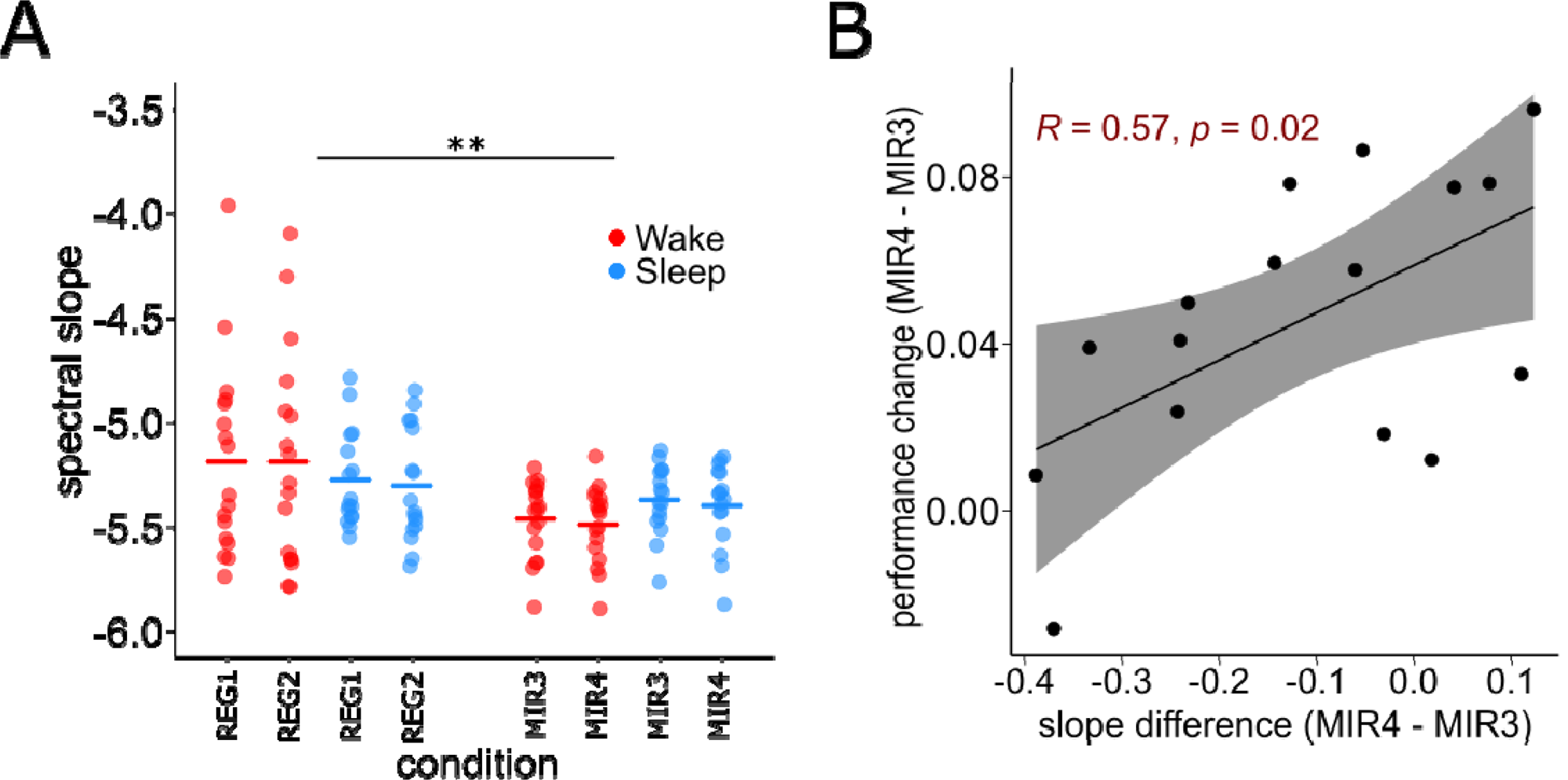
Spectral slope tracks motor adaptation and predicts sleep-mediated improvement in motor adaptation. A) Slope of the power spectral density averaged over all EEG electrodes, steeper during mirrored typing as compared to regular typing. B) Change in the spectral slope from the pre- to post-sleep mirrored typing sessions correlated with the change in performance in the mirrored typing session from pre- to post-sleep (MIR4 – MIR3). Note that this correlation was not significant in the wake group (Fig. S7H). Each point in panel A represents the mean over one subject and the crossbars represent the mean over all subjects. REG1: Regular typing before retention, REG2: Regular typing after retention, MIR3: Mirrored typing before retention, MIR4: Mirrored typing after retention. R = Pearson’s correlation coefficient. ** p < 0.001.

**Table 2.**
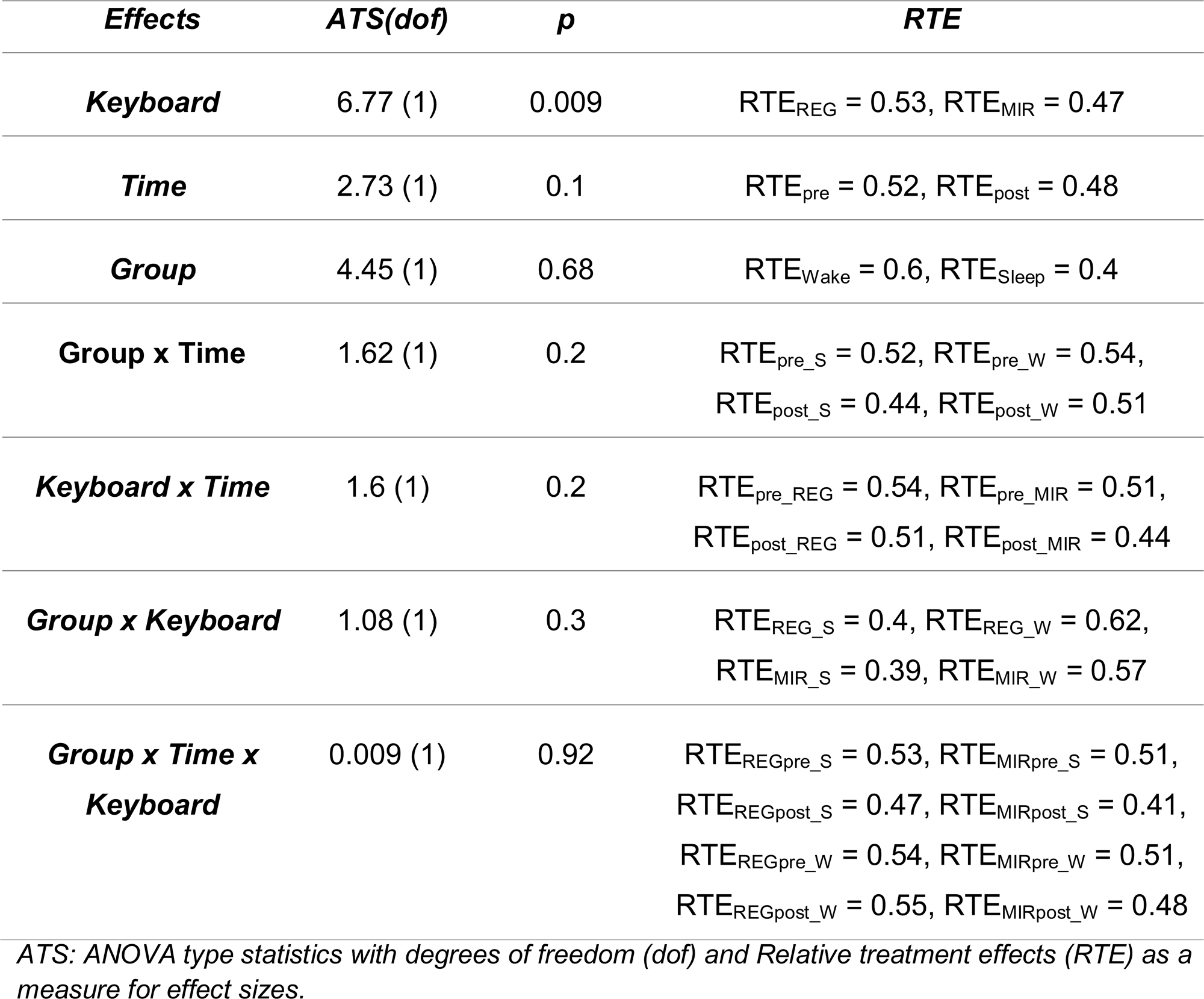
Statistical results for the difference in the spectral slope.

## 4. Discussion

In this study, we employed a novel motor adaptation paradigm, capitalizing on the extensive training that participants received on typing on computer keyboards throughout their lives, to leverage a fine-motor adaptation task where participants had to adapt to typing on a new, vertically mirrored keyboard (Fig. 1). We measured the typing performance on the regular and the vertically mirrored keyboard before and after a retention period that included either 8h of nocturnal sleep or a similar period of daytime wakefulness.

Our results demonstrate that post-learning sleep but not wakefulness was beneficial to typing on the mirrored keyboard (Fig 2A). After nocturnal sleep indeed, participants attained performance levels achieved through training, whereas it declined after wakefulness (Fig 2B). The effects of post-learning sleep, however, were limited to the early phase of the testing session, as performance in the wake group then increased to reach the same level than the other group, suggesting a transient nature of the impact of sleep on motor adaptation, at least as compared to the amplitude of actual training effects. Importantly, the beneficial effects of sleep on motor adaptation did not interfere with the already automated skill of typing on a regular keyboard (Fig. S2).

Our EEG results demonstrate a change in typing-related brain activity following nocturnal sleep (Fig 3). Already before the retention interval, there was a decrease in central beta (13-30Hz) power during mirrored typing as compared to regular typing (Fig. 3A; Fig. S3A). Furthermore, there was a general decrease in beta power after sleep, observed both during mirrored and regular typing trials (Fig. 3A). These results suggest a general modulation of beta power by post-training sleep. To rule out an effect of circadian variation, we compared beta power during resting state recording in the evening prior to training and in the morning prior to testing (Fig. S3B). Beta power actually increased after a night of sleep, indicating that the observed decrease in beta power following sleep is specifically related to the motor adaptation task, rather than a circadian effect.

Using multivariate decoding, beta power exhibited the highest classification accuracy in distinguishing between regular and mirrored typing trials (Fig. S3A). Crucially, however, we observed a significant decline in the classification accuracy in the beta band following sleep (Fig. 3B) indicating an increased similarity/less differentiation in brain activity between the regular and mirrored typing tasks only after sleep. These findings suggest that the role of sleep in consolidating motor adaptation skills might in part rely on integrating the movement representations of the mirrored typing task into the well-established networks associated with the automatized regular typing task (Bastian, 2008). Additionally, as beta power during regular typing also decreased after sleep (Fig. 3A), it could be surmised that the positive influence of sleep is not specific to the adaptation process, but also benefits de-adaptation, i.e., going back to the automatized process (regular typing) after adaptation training (Davidson & Wolpert, 2004).

During sleep, although we were not able to confirm our initial hypothesis with role of REM sleep and REM beta oscillations in motor adaptation (Fig. S6), we demonstrated that the improvement in motor adaptation performance correlated with the increase in fast spindle density mainly in N2 sleep. This result confirms previous findings that linked N2 fast spindles with the consolidation of motor memories (King et al., 2017). We could not observe this correlation during N3. This might be because during N3 sleep the consolidation relies on the coupling of spindles of fast spindles with slow oscillations and not necessarily on the increase in fast spindle density (Thürer et al., 2018). Previous research suggests that spindles are modulated over learning-related areas (Cox et al., 2014; Petzka et al., 2022). The fact that we observed this correlation over 10 electrodes (similar to global beta effects; Fig. S3A) might indicate the high cognitive demand for adaptation in the brain.

In addition to the changes in periodic EEG activity, the spectral slope of the EEG signal, a measure of aperiodic brain activity, became steeper during motor adaptation (mirrored typing) as compared to the original task (regular typing) (Fig. 5A). This indicates increased inhibitory activity during adaptation. Importantly, after sleep, we observed a flattening of the slope during mirrored typing trials, correlated with improved adaptive performance (Fig. 5B). No such correlation was observed after wakefulness (Fig. S4G). These findings suggest that sleep facilitates motor adaptation by influencing cortical excitation-inhibition balance (Bridi et al., 2020) and provides additional support for the advantageous role of sleep in modulating cortical inhibition during motor adaptation.

To the best of our knowledge, this study presents novel findings demonstrating an interaction between sleep, periodic, and aperiodic brain activity which benefits motor adaptation performance. Our results align with the concept that beta band activity plays a role in maintaining the status quo in the brain and constraints flexibility (Engel and Fries, 2010). By demonstrating an influence of post-training sleep on beta power, we extend previous results showing that a decrease in beta power is indicative of adaptive motor behaviour in the brain (Darch et al., 2020)

What are the underlying mechanisms of motor adaptation? Previous research has highlighted the association between beta oscillation changes and variations in the activity of GABAergic interneurons (Jensen et al., 2005; Rossiter et al., 2014). Jensen et al. (2005) demonstrated a correlation between increased GABA-mediated inhibition and enhanced beta oscillatory power. Thus, it is plausible that the reduction in the beta power reflects the release of cortical inhibition, i.e., dis-inhibition. This dis-inhibition may be advantageous in dealing with the heightened cognitive demands associated with motor adaptation.

The notion of sleep-mediated dis-inhibition that benefits (de-)adaptation was further examined by measuring the difference in the slope of the EEG signal during regular and mirrored typing sessions. Previous research by Gao et al. (2017) using a computational model of cortical circuits demonstrated that a steeper spectral slope corresponds to increased recruitment of inhibitory populations, while a flatter slope reflects reduced inhibition. In this line of reasoning, a steeper slope during initial motor adaptation training and a correlation between the flattening of the slope and improved performance provide supportive evidence for a sleep-induced cortical disinhibition that promotes motor adaptation. Cortical inhibition has been shown to be crucial for continual learning as well as for protecting overlapping memories from interfering with each other (Barron, 2021). Following the initial learning phase, sleep promotes such inhibition in the cortex to facilitate memory consolidation (Bridi et al., 2020; Niethard et al., 2017). Our results suggest that this inhibition might also be mediated by fast sleep spindles activity during N2 sleep. Indeed, sleep spindles have been shown to promote inhibition in the cortex (for a review see: Fernandez and Luthi, 2019) and increase in number after motor adaptation training (Solano, 2022). In the post-sleep testing session, hippocampus-mediated dis-inhibition of neocortical circuits might have facilitated memory recall, contributing to the improvement of motor adaptation (Koolschijn et al., 2020). According to such a model of cortical dynamics, the initial increase in inhibition, suggested by decreased beta power and a steeper slope during motor adaptation training, would facilitate memory encoding and protect against interference from overlapping memories (regular typing). Subsequently during sleep, fast sleep spindles would support the consolidation process by regulating the balance of excitation and inhibition. Finally, during recall, the release of inhibition would become crucial for efficient switching between different motor networks that govern various typing strategies.

Altogether, our findings shed light on the potential use of periodic (beta power) and aperiodic (spectral slope) EEG parameters as markers of motor adaptation. Additionally, our results provide insights into the regulatory role of sleep spindles in facilitating motor adaptation. This model aligns with recent theories of memory consolidation, proposing that the interplay between periodic activity and aperiodic activity plays a vital role in memory consolidation processes (Helfrich et al., 2021). Future investigations should aim to discern the specific contributions of periodic and aperiodic activity to sleep-related memory consolidation processes.

Our study has a few limitations. First, we cannot rule out the presence of a motor-learning component in our task. Participants had to perform a series of five-finger movements to type six (MIR1-4) or 12 (MIR5) different words on the mirrored keyboard and therefore, they might have learned to perform a motor sequence of six finger movements to type the words, more than adapting their typing behaviour according to previous knowledge. However, since we recruited experts in touch-typing, we assume that participants must first inhibit a specific and very strong motor plan before being able to perform the new adaptation task. Therefore, even if the task is somewhat contaminated by traces of motor learning activity, the prevailing behaviour would still be the adaptation of fine-motor skills. In the same vein, we did not control for the ability of the participants to switch between different keyboard setups (e.g., QWERTZ - QWERTY) which might entail some experience with motor adaptation. However, as the task specifically tackles *mirrored* typing, which is highly uncommon, we assume that experience with usual keyboards switching would be irrelevant for our task. Another potential limitation is a possible circadian influence on beta power, and the difference in baseline beta power between the sleep and wake groups despite a random assignment of participants to these conditions. It makes it possible that time of day differences between training and testing may have had an impact on beta power values. A previous study showed that beta activity during pre-movement desynchronization periods increased from morning to afternoon, while it remained steady throughout the day during post-movement rebound periods (Wilson et al., 2014). However, our control analysis demonstrated that decreased beta power between the pre- and the post-sleep sessions was task related, since resting-state beta power increased from the eyes-closed resting states before training to the one before testing(Fig. S3C), i.e., the opposite pattern to the task-related beta power. Moreover, there was no significant change in beta power in the wake group, corroborating the notion that decreased beta power was modulated by post-training sleep, and not just a mere circadian effect. On another aspect, the all-male sample recruited for this study is of course a limitation when it comes to the generalisability of the results to the whole population. Future studies should elaborate on gender differences in the performance and consolidation in fine-motor skill tasks.

Finally, it must be mentioned that we deviated from the pre-registered analyses by adding an exploratory analysis of the EEG slope (Fig. 5). This analysis was motivated by the growing evidence linking the EEG spectral slope with changes in cognitive functions (Waschke et al.,2021; Hohn et al., 2022), and consciousness levels (Lendner et al., 2020). This exploratory analysis in our study provides more information regarding the dynamics of excitation and inhibition in the cortex and their role in mediating adaptive processes in the brain.

In summary, our findings provide evidence for the involvement of sleep in the adaptation of highly automated fine-motor skills. We observed significant changes in cortical oscillatory and aperiodic activity following nocturnal sleep, which parallel the positive influence of sleep on motor adaptation. Our findings suggest that sleep exerts a beneficial impact on motor adaptation by facilitating the context-specific release of cortical inhibition. Additionally, sleep facilitates the integration of newly acquired adaptation tasks into pre-existing motor memory networks while safeguarding against interference between these overlapping memory traces. The underlying mechanisms through which sleep promotes cortical disinhibition and protects against memory interference warrant further investigation in future studies.

## Acknowledgements

This work was supported by the Austrian Science Funds (FWF) and the Austrian Academy of Sciences (OEAW).

---

**Figure S1.**
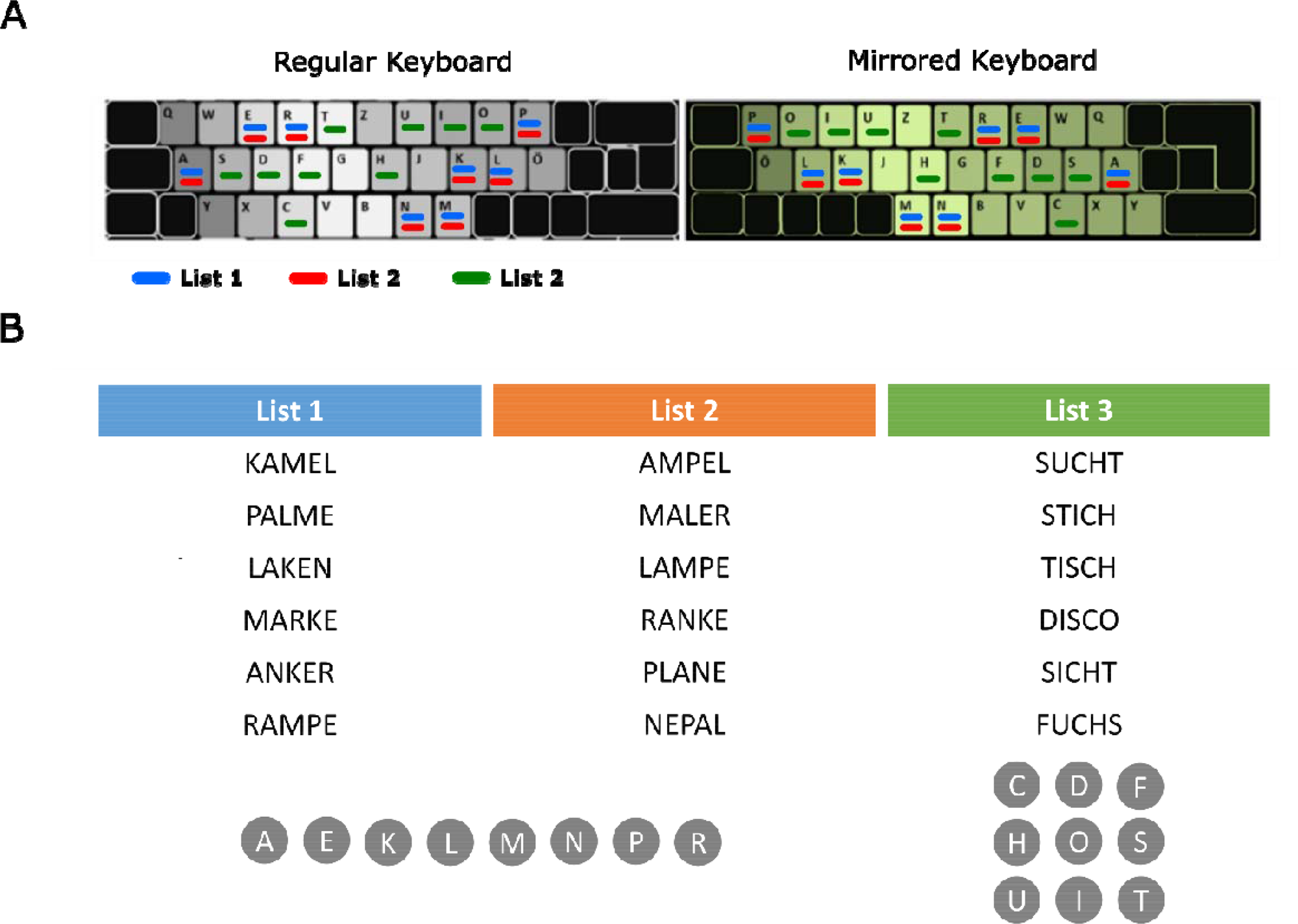
The mirrored keyboard and the wordlists. A) (Left) a picture of a regular German layout keyboard. (Right) a picture of the new, mirrored German layout keyboard that was used for motor adaptation. Note that we did not physically change the buttons on the mirrored keyboard as depicted in the picture. Instead, we re-programmed the commands of the keyboard so that the buttons typed the mirrored letters. B) The different wordlists we used for this experiment. Note that Lists 1 and 2 contained different words that consisted of the same letters. List 3 contained new words that consisted of a new set of letters. Also note that the words in List 1 were the only words that the participant typed on the mirrored keyboard during the training session before the retention interval.

**Figure S2.**
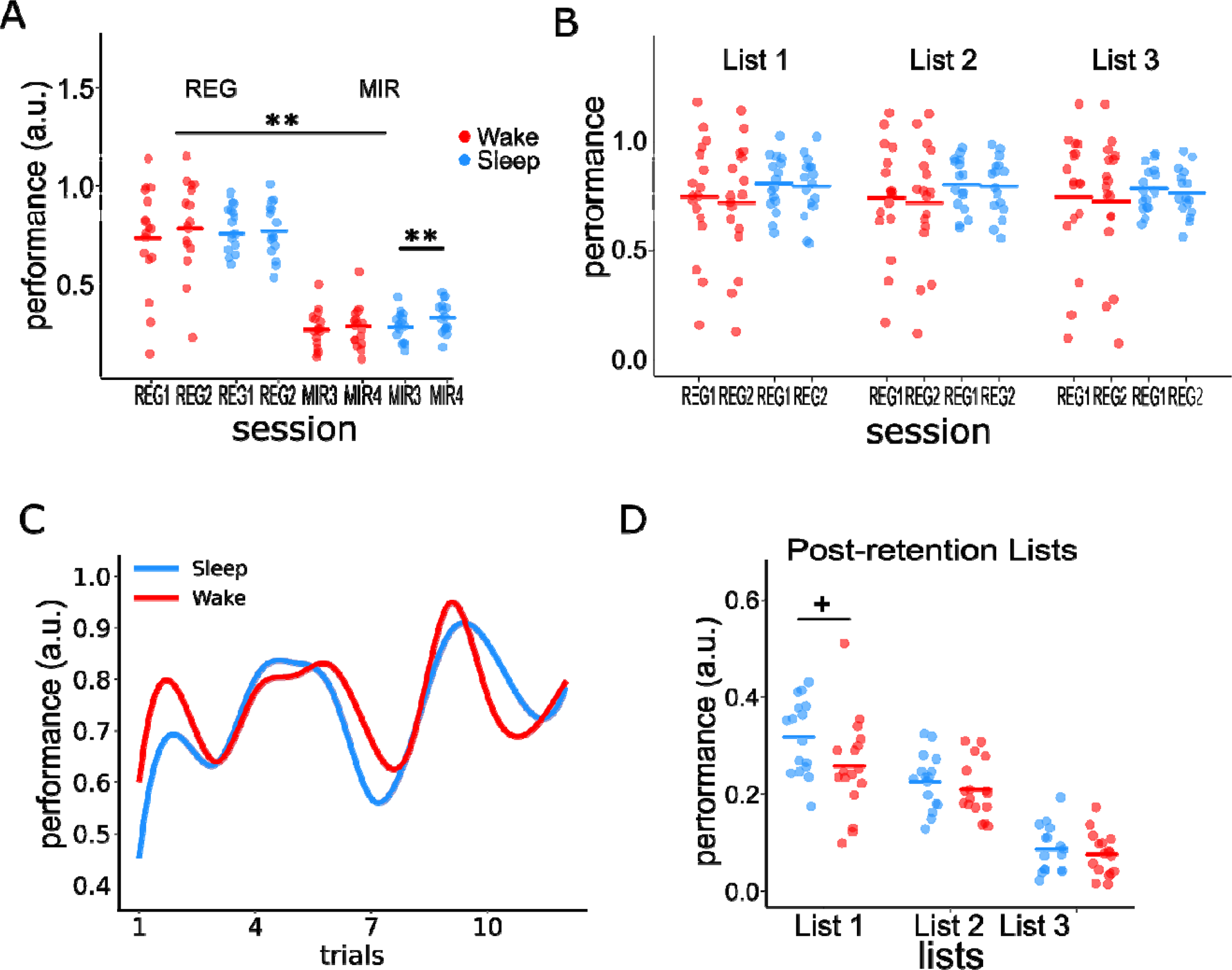
Extended behavioural results. A) Behavioural performance during regular and mirrored typing sessions (an extended analysis of Fig. 1C). A 2×2x2 mixed non-parametric test with within-subject factors Keyboard (REG vs. MIR) and Time (before vs. after retention), and a between-subjects factor Group (sleep vs. wake). The results showed main effects of Keyboard (*ATS(1) = 220.77, p < 0.001, RTE_REG_ = 0.73, RTE_MIR_ = 0.27*), and Time (*ATS(1) = 8.67, p = 0.003, RTE_pre_ = 0.49, RTE_post_ = 0.51*). Typing performance was higher during the REG than the MIR typing trials as well as after retention than before retention. There was no difference in REG and MIR performance between sleep and wake group (*ATS(1) = 0.92, p = 0.34, RTE_S_ = 0.47, RTE_W_ = 0.52*). Further, we found a significant interaction for the factors Keyboard x Time (*ATS(1) = 18.2, p_bonf_ < 0.001, RTE_REGpre_ = 0.72, RTE_MIRpre_ = 0.25, RTE_REGpost_ = 0.73, RTE_MIRpost_ = 0.29*). Post-hoc tests revealed that performance on the mirrored keyboard increased after the retention (*ATS(1) = 18.41, p_bonf_ < 0.001, RTE_pre_ = 0.45, RTE_post_ = 0.55*), while performance on the regular keyboard remained unchanged (*ATS(1) = 0.784, p_bonf_ = 0.75, RTE_pre_ = 0.49, RTE_post_ = 0.51*). Finally, we observed a three-way interaction, Keyboard x Time x Group (*ATS(1) = 12.82, p < 0.001, RTE_REGpre_S_ = 0.75, RTE_MIRpre_S_ = 0.27, RTE_REGpost_S_ = 0.74, RTE_MIRpost_S_ = 0.35, RTE_REGpre_W_ = 0.7, RTE_MIRpre_W_ = 0.23,, RTE_REGpost_W_ = 0.71, RTE_MIRpost_W_ = 0.25*). Post-hoc test showed that typing performance on the mirrored keyboard improved after sleep (*ATS(1) = 25.27, p_bonf_ < 0.001, RTE_pre_ = 0.42, RTE_post_ = 0.58*) but not after wakefulness (*ATS(1) = 1.63, p_bonf_ = 0.81, RTE_pre_ = 0.48, RTE_post_ = 0.52*) thus confirming our hypothesis that sleep benefits motor adaptation consolidation. There was no change in the typing performance on the regular keyboard after retention neither in the sleep group (*ATS(1) = 1.26, p_bonf_ = 0.26, RTE_pre_ = 0.51, RTE_post_ = 0.49*) nor in the wake group (*ATS(1) = 3.65, p_bonf_ = 0.22, RTE_pre_ = 0.48, RTE_post_ = 0.52*). B) Sleep’s positive influence on motor adaptation did not interfere with the original task of regular typing. B) Typing performance on the regular keyboard in the different lists (List 1-3) before and after retention. Note that we used only words in List 1 for the training on the mirrored keyboard before retention; words in List 2 and List 3 were typed only on the regular keyboard before retention. A 2×2X2 mixed non-parametric test with two within-subject factors; *Lists* (List1, 2, and 3) and *Time* (before vs. after retention), and one between-subjects factor; *Group* (Sleep vs. Wake) showed only a main effect of *TIME* (*ATS(1) = 24.612, p < 0.001, RTEpre = 0.51, RTE_post_ = 0.49*) as performance increased after retention. There were no significant differences between the Lists (*ATS(1.63) = 0.267, p = 0.719, RTE_List1_ = 0.51, RTE_List2_ = 0.5, RTE_List3_ = 0.49*), or *Group* (*ATS(1) = 0.064, p = 0.801, RTE_Sleep_ = 0.49, RTE_wake_ = 0.51*). There was a significant interaction *Group X List* (*ATS(1) = 0.064, p = 0.801, RTE_Sleep_ = 0.49, RTE_wake_ = 0.51*). There was no significant interaction *Time X Lists X Group* (*ATS(1.99) = 0.495, p = 0.609*). C) No interference effects of sleep on regular typing. We compared the performance between sleep and the wake group on the words of List 1 (which participants trained on) before the retention on a word-by-word level. There was no significant difference in the performance between the groups (P>0.05). D) Comparisons between the different word lists (Lists 1, 2 and 3) after retention. Note that List 1 words were presented in MIR4, while MIR5 session contained words from lists 2 and 3. Results showed no effect of Group (*ATS(1) = 3.725, p = 0.122, RTE_sleep_ = 0.54, RTE_wake_ = 0.46*), but a main effect of *List* (*ATS(1.943) = 51.237, p < 0.001, RTE_List1_ = 0.74, RTE_List2_ = 0.58, RTE_List3_ = 0.18*). Performance was significantly higher in List 1 than List 2 (*ATS(1) = 62.719, p_bonf_ < 0.001, RTE_List1_ = 0.62, RTE_List2_ = 0.38*) and List 3 (*ATS(1) = 23.537, p < 0.001, RTE_List1_ = 0.74, RTE_List3_ = 0.26*), and List 2 than List 3 (*ATS(1) = 196.535, p < 0.001, RTE_List2_ = 0.74, RTE_List3_ = 0.26*). Finally, there was a significant interaction *Group X List* (*ATS(1.943) = 80.535, p = 0.031, RTE_List1_sleep_ = 0.82, RTE_List2_sleep_ = 06, RTE_List3_sleep_ = 0.2, RTE_List1_wake_ = 0.66, RTE_List2_wake_ = 0.56, RTE_List3_wake_ = 0.17*). Post-hoc Wilcoxon rank sum tests showed that for List 1 the performance was marginally higher for the sleep group as compared to the wake group (*W = 212, p_bonf_ = 0.081, 95% confidence interval [0.006 0.13]).* Each point in panel A represents the mean over one subject and the crossbars represent the mean over all subjects. REG1: Regular typing before retention, REG2: Regular typing after retention. MIR3: Mirrored typing before retention, MIR4: Mirrored typing after retention.

**Figure S3.**
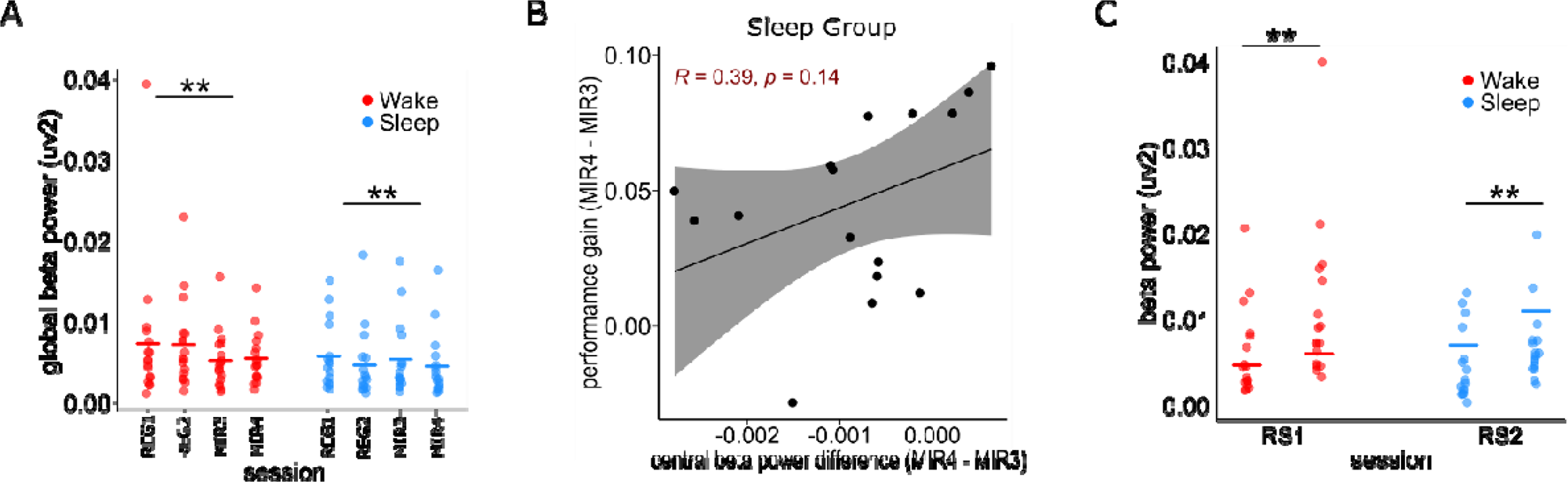
Spatial and temporal aspects of movement-related beta band activity. A) Global beta power differences between regular and mirrored typing. A 2×2x2 mixed non-parametric test with the two within-subjects Keyboard (REG vs. MIR) and Time (pre-retention vs. post-retention) and a between-subject factor Group (Sleep vs. Wake) showed that beta power was lower during mirrored typing (*ATS(1) = 13.57, p < 0.001, RTE_REG_ = 0.52, RTE_MIR_ = 0.48*). We found no effect of Time (*ATS(1) = 1.39, p = 0.23, RTE_pre_ = 0.51, RTE_post_ = 0.49) or Group (ATS(1) = 1.24, p = 0.27, RTE_Wake_ = 0.55, RTE_Sleep_ = 0.44*). However, there was a significant interaction Group x Time (*ATS(1) = 20.86, p < 0.001, RTE_SLpre_ = 0.49, RTE_SLpost_ = 0.4, RTE_Wpre_ = 0.53, RTE_Wpost_ = 0.58*), as beta power decreased after sleep (*ATS(1) = 20.29, p < 0.001, RTE_SLpre_ = 0.49, RTE_SLpost_ = 0.4*) but increased after wakefulness (*ATS(1)= 7.815, p = 0.01, RTE_SLpre_ = 0.49, RTE_SLpost_ = 0.4*). There was no interaction *Keyboard x Time* (*ATS(1) = 0.77, p = 0.38*). We also observed a significant interaction Keyboard x Time x Group *(ATS(1) = 4.33, p = 0.04, RTE_REGpre_S_ = 0.51, RTE_MIRpre_S_ = 0.46, RTE_REGpost_S_ = 0.4, RTE_MIRpost_S_ = 0.4, RTE_REGpre_W_ = 0.55, RTE_MIRpre_W_ = 0.5,, RTE_REGpost_W_ = 0.62, RTE_MIRpost_W_ = 0.54*). Post-hoc tests showed that beta power decreased after sleep during both regular (*ATS(1) = 18.65, p < 0.001, RTE_pre_ = 0.56, RTE_post_ = 0.44*) and mirrored typing (*ATS(1) = 9.04, p = 0.01, RTE_pre_ = 0.54, RTE_post_ = 0.46*) but did not significantly differ from pre to post-wakefulness (Regular: *ATS(1) = 5.639, p = 0.07, RTE_pre_ = 0.47, RTE_post_ = 0.53; Mirrored: ATS(1) = 2.069, p = 0.6, RTE_pre_ = 0.48, RTE_post_ = 0.52*). B) The correlation between the change in beta power after sleep with the motor adaptation performance change from pre- to post-sleep. The change in beta power from pre-sleep to post-sleep mirrored typing session did not correlate with the change in behavioural typing performance on the mirrored keyboard. C) The effects of sleep on beta power are task related. A 2×2 mixed non-parametric test with Time as within subject variable and Group as between subject variable Comparing beta power during the eyes-closed resting state period before training (RS1) and before testing (RS2) revealed no difference in beta power between groups (*ATS(1) = 1.51, p = 0.22, RTE_Wake_ = 0.55, RTE_Sleep_ = 0.45*). However, there was a time effect as beta power increased from pre-retention to post-retention in both groups in the opposite direction of the task-related beta changes after sleep (*ATS(1) = 24.82, p < 0.001, RTE_SLpre_ = 0.35, RTE_SLpost_ = 0.54, RTE_Wpre_ = 0.42, RTE_Wpost_ = 0.68*). There was no significant interaction *Time X Group* (*ATS(1) = 0.52, p = 0.47*)

**Figure S3.**
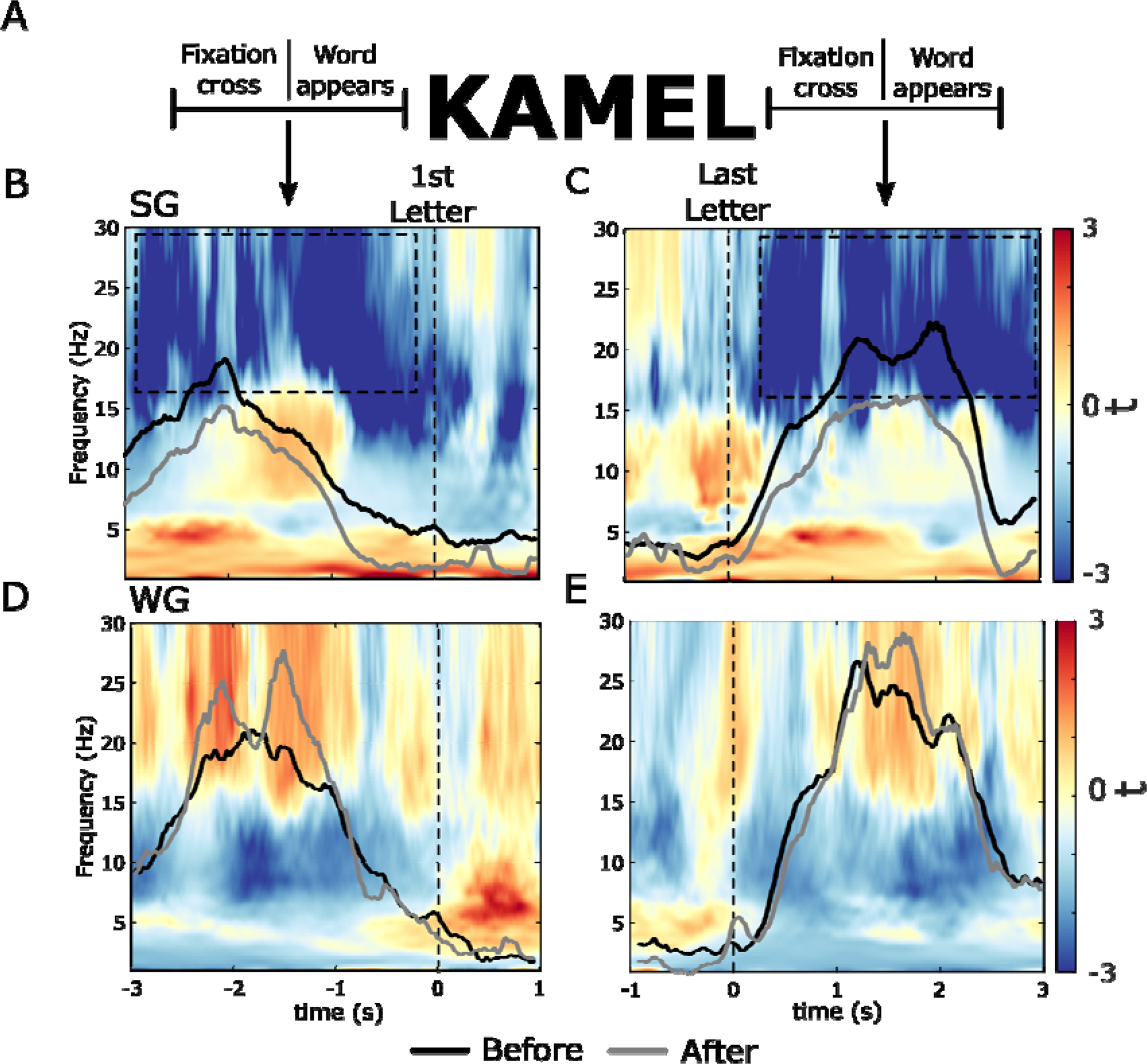
Temporal dynamics of beta power during mirrored typing trials. A) As participants had to type five letter words, to investigate pre-movement beta desynchronization and post-movement beta rebound periods, we looked at three second windows before typing the first letter of every word, and the three seconds following typing the last letter of each word, respectively. Note that in both cases, the analysed time window included the display of the fixation cross (1.5s) either before or after typing (for pre-movement and post-movement, respectively), and the appearance of a word on the screen, but did not include any typing activity. To confirm that there was no typing activity in this period, we measured the typing latency per session starting from the display of the fixation cross until typing the first letter of the word (desynchronization time) as well as the average time after typing the last letter of the word until typing the first letter of the following word (rebound window). The mean desynchronization time was 3.06s ± 0.48s (mean ± standard deviation) during MIR3 and 2.86s ± 0.69 during MIR4, while the mean rebound time was 4.17s ± 0.33 during MIR3 and 3.81s ± 0.38 during MIR4. B) Time-Frequency representations demonstrated a general decrease in the power of beta band following sleep during the pre-movement beta desynchronization periods (∑t(15) = -14006.95, p = 0.002) as well as (C) during post-movement beta rebound periods (t(15) = -18395.25, p < 0.001). We did not observe this decrease in beta power following a period of wakefulness (D-E). Note the black and grey lines on top of the time-frequency plots that indicate beta power averaged over C3 and C4, showing less beta desynchronization after sleep but not after wakefulness and lower rebound after sleep but not after wakefulness. The difference, however, was not statistically significant. REG1: Regular typing before retention, REG2: Regular typing after retention. MIR3: Mirrored typing before retention, MIR4: Mirrored typing after retention. SG: Sleep group, WG: Wake group.

**Figure S5.**
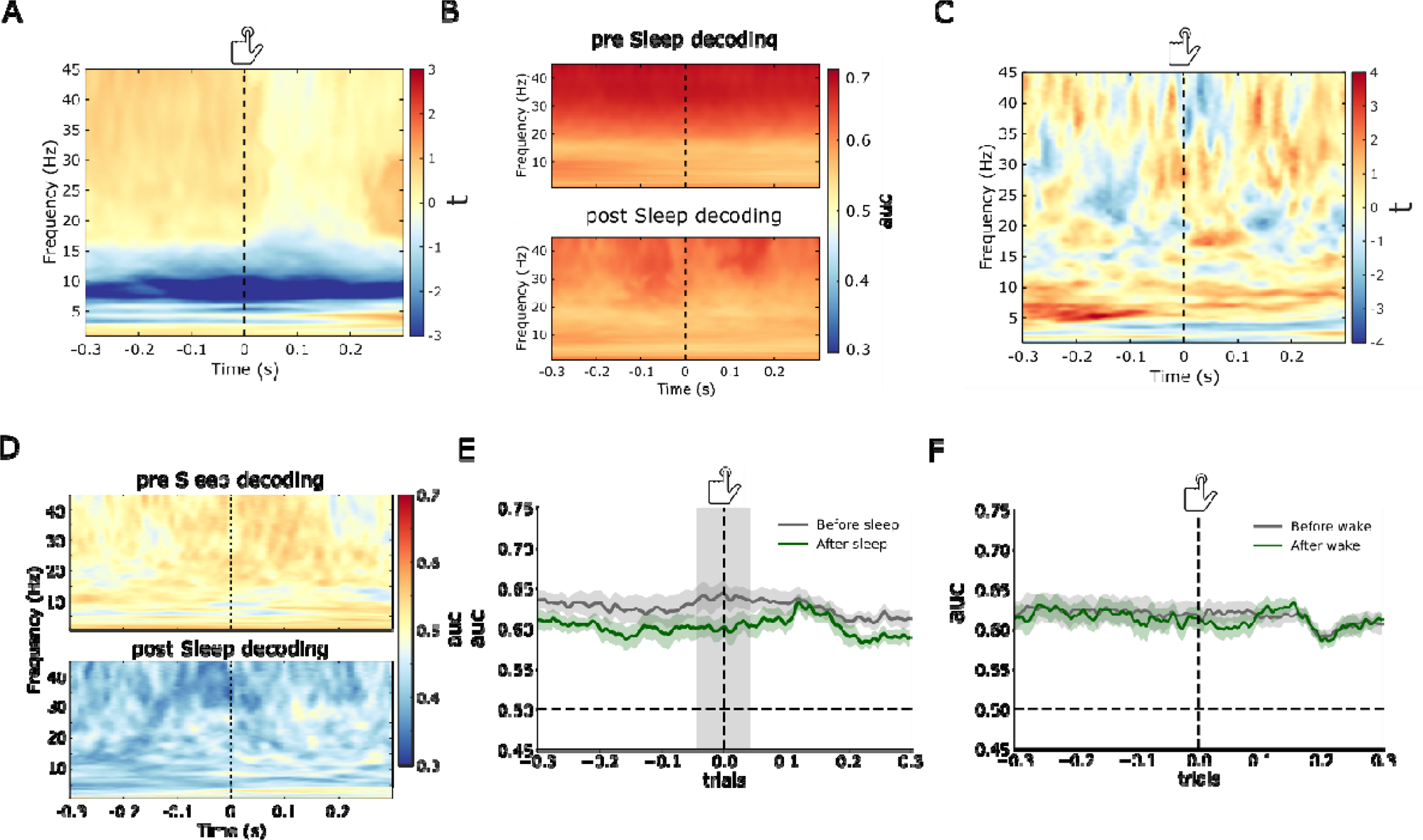
Decoding results between regular vs. mirrored typing and correct vs. incorrect mirrored typing trials. A) Frequency decoding on central electrodes in the wake group showed no change in beta band decoding accuracy (area under curve; auc) after retention. There was a decrease in decoding performance after wakefulness in the theta/alpha range that was, however, insignificant. B) Frequency decoding on central electrodes before and after sleep. Time-frequency decoding accuracy before and after sleep to show that beta band demonstrated the highest decoding performance before and after sleep. Note the decrease in decoding performance in the beta range after sleep. C) Frequency decoding of correct vs. incorrect mirrored typing trials in the wake group did not change from before to after retention (Post-wake – Pre-wake). D) A time-frequency representation of the frequency decoding on correct vs incorrect trials before and after sleep. E-F) According to our registered analysis plan, we decoded regular vs. mirrored typing using time-locked data averaged over all electrodes in the -0.3s to 0.3s window. E) Decoding performance dropped significantly after sleep (grey shading) in the time window *(-0.04s – 0.02s;* ∑*t(15) = -112.776, p = 0.012)*, but not after wakefulness (F).

**Figure S6.**
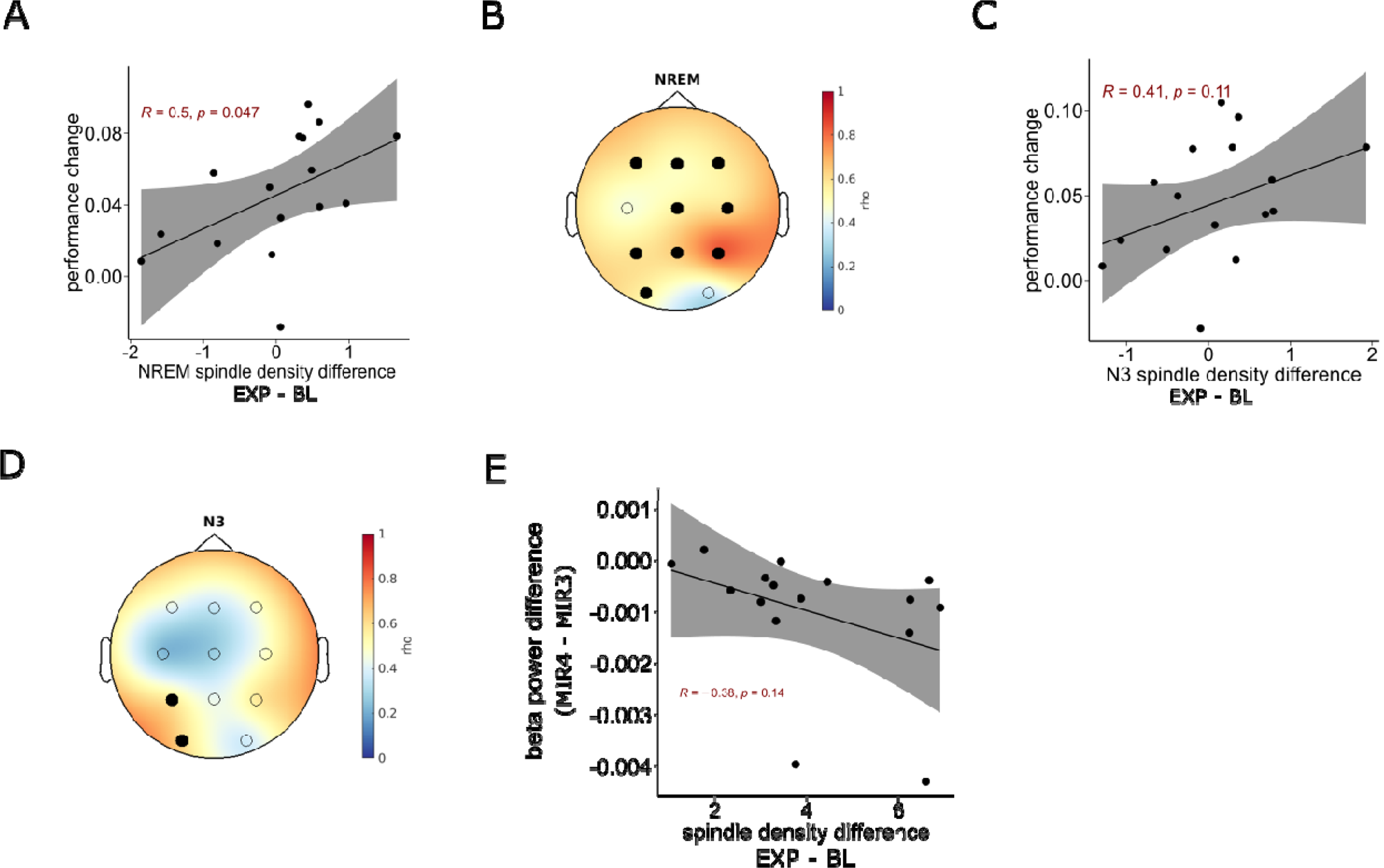
Sleep results. A-D) Fast spindle density (averaged over C3 and C4) correlation with the change in mirrored typing performance after sleep. A) When we measured fast sleep spindle density over the whole duration of N2 and N3 sleep combined we observed a significant positive correlation (*r(14) = 0.5, p = 0.047*). B) The topography of correlation resembled that observed during N2 indicating a significant cluster of 9 electrodes (All except C3 and O2; ∑*t(15) = 26.35, p = 0.003, d = 0.32*). C) N3 Fast sleep spindles density showed no significant correlation with the change in performance (*r(14) = 0.41, p = 0.11*). D) Topography of electrodes exhibiting significant correlation of the change in fast spindle density with the change in mirrored typing performance (P1 and O1; ∑*t(15) = 5.51, p = 0.03, d = 0.34*). E) The change in NREM stage 2 fast sleep spindle density from the baseline night to the experimental night did not correlate with the change in beta power from pre- to post-sleep during mirrored typing sessions. REM: rapid eye movement sleep. R: Pearson’s correlation coefficient. Rho: Spearman’s correlation. BL: baseline night, EXP: experimental night.

**Figure S7.**
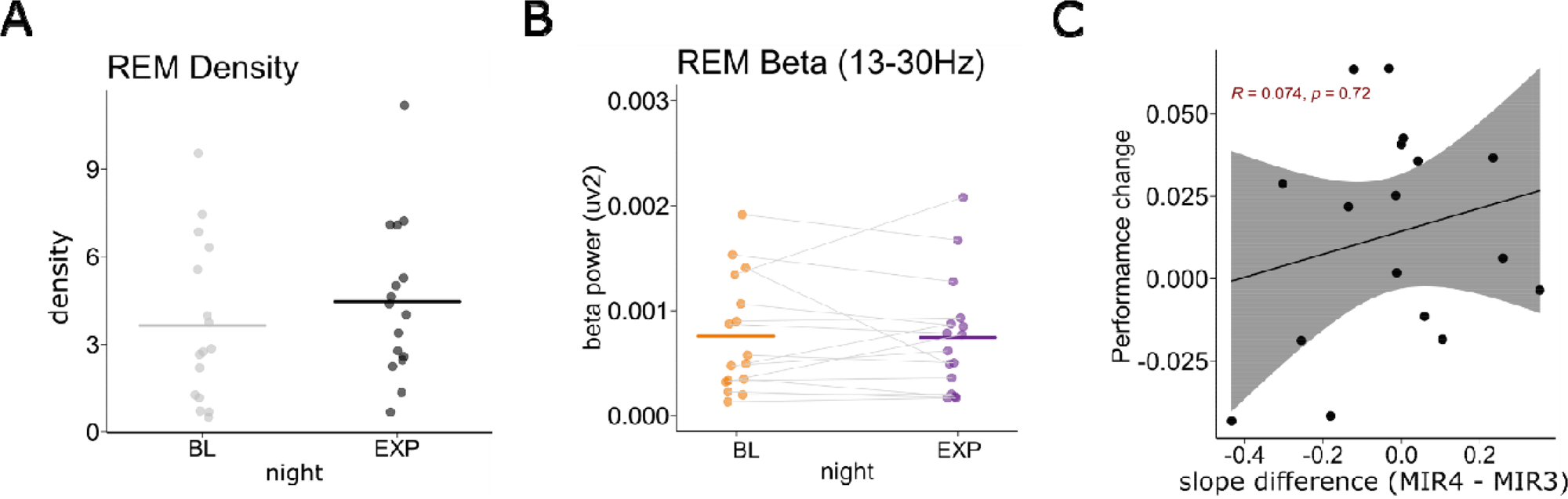
REM beta power did not differ between baseline and experimental nights. According to our registered analysis, we compared REM activity during the baseline night and the experimental night. A) No significant differences in REM sleep density (time spend in REM / Total sleep time) between the baseline night and the experimental night (ATS(1) = 1.36, p = 0.24, RTEBL = 0.55, RTE_Exp_ = 0.44). B) Beta power during REM sleep did not change from the baseline to the experimental night (ATS(1) = 0.03, p = 0.87, RTE_BL_ = 0.5, RTE_Exp_ = 0.5). C) Spectral slope correlation with performance in the wake group. The change in slope from pre- to post-wakefulness did not correlate with the change in mirrored typing performance (MIR4 - MIR3).

