## Supplemental Figures document for "Post-training sleep modulates motor adaptation and task-related beta oscillations"

6.
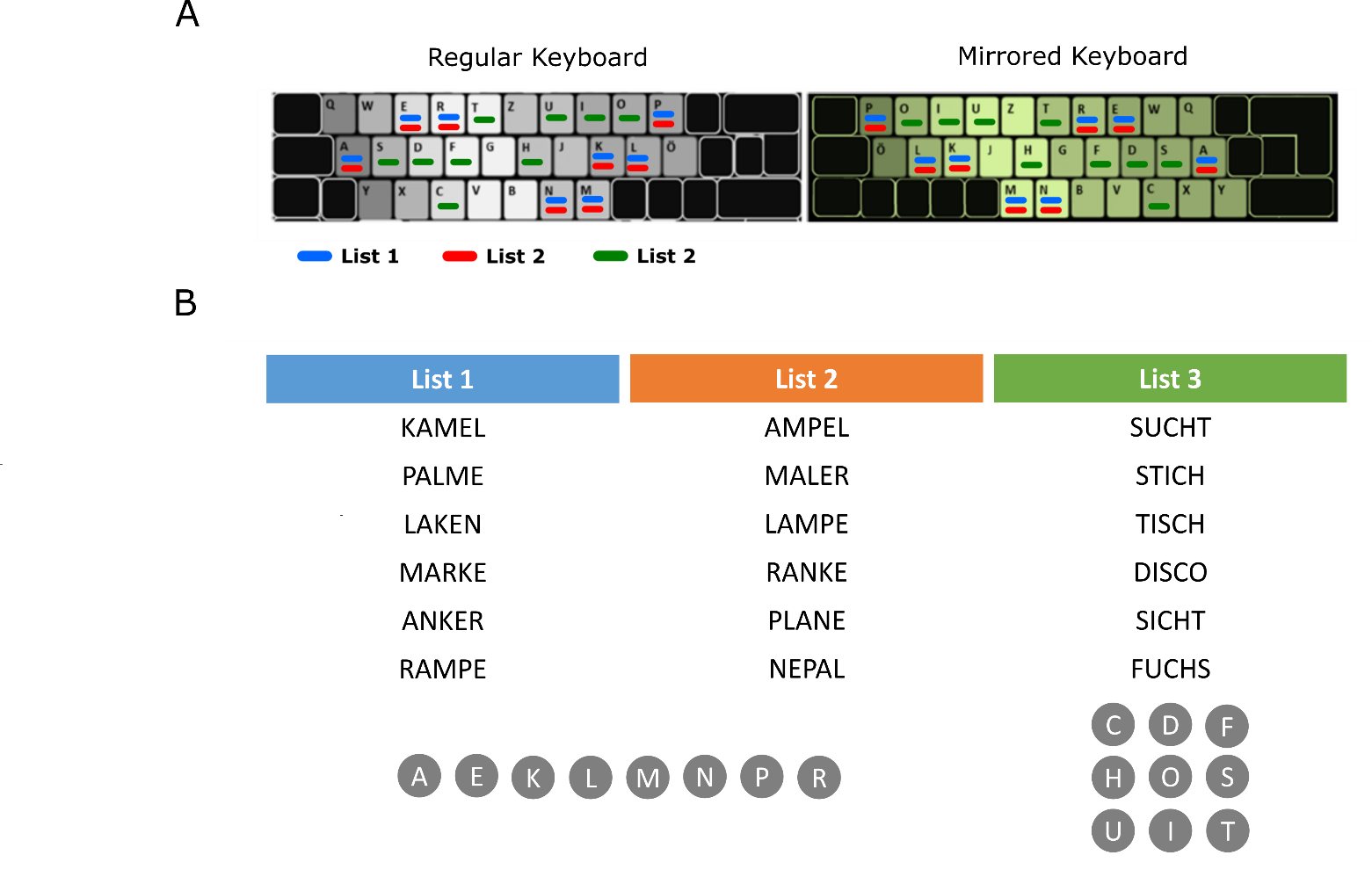
**Supplementary material legends**

**Figure S1. The mirrored keyboard and the wordlists.** A) (Left) a picture of a regular German layout keyboard. (Right) a picture of the mirrored German layout keyboard that was used. Note that we did not physically change the buttons on the mirrored keyboard as depicted in the picture. Instead, we re-programmed the commands of the keyboard so that the buttons would type the mirrored letters. B) The different wordlists we used for this experiment. Note that Lists 1 and 2 contained different words that consisted of the same letters. List 3 contained new words that consisted of a new set of letters. Also note that the words in List 1 were the only words that the participant typed on the mirrored keyboard during the training session before the retention interval.

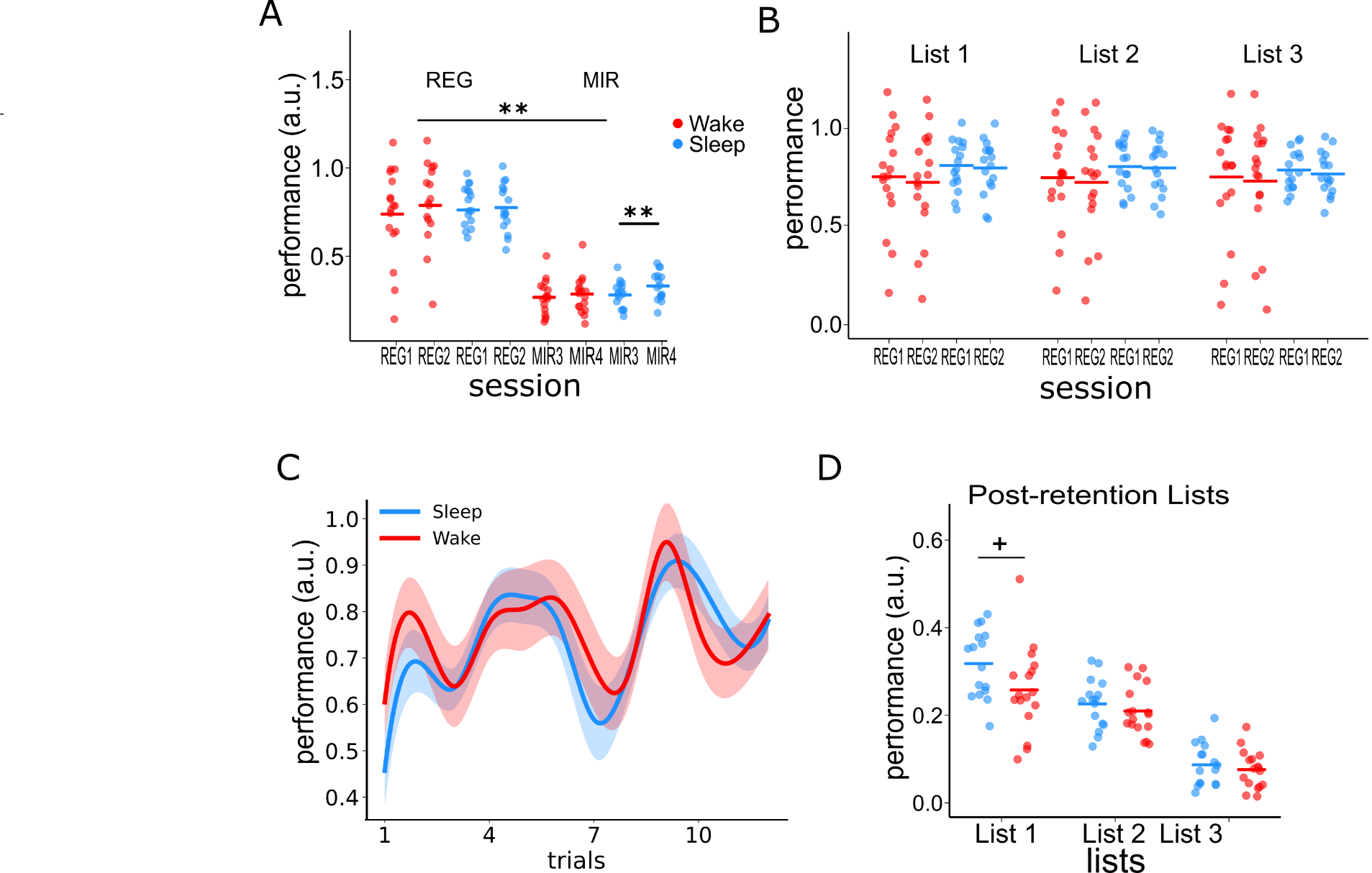

**Figure S2. Extended behavioural results.** A) Behavioural performance during regular and mirrored typing sessions (an extended analysis of Fig. 1C). A 2x2x2 mixed non-parametric test with within-subject factors Keyboard (REG vs. MIR) and Time (before vs. after retention), and a between-subjects factor Group (sleep vs. wake). The results showed main effects of Keyboard (*ATS(1) = 220.77, p < 0.001, RTE_REG_ = 0.73, RTE_MIR_ = 0.27*), and Time (*ATS(1) = 8.67 , p = 0.003, RTE_pre_ = 0.49, RTE_post_ = 0.51*). Typing performance was higher during the REG than the MIR typing trials as well as after retention than before retention. There was no difference in REG and MIR performance between sleep and wake group (*ATS(1) = 0.92, p = 0.34, RTE_S_ = 0.47, RTE_W_ = 0.52*). Further, we found a significant interaction for the factors Keyboard x Time (*ATS(1) = 18.2, p_bonf_ < 0.001, RTE_REGpre_ = 0.72, RTE_MIRpre_ = 0.25, RTE_REGpost_ = 0.73, RTE_MIRpost_ = 0.29*). Post-hoc tests revealed that performance on the mirrored keyboard increased after the retention (*ATS(1) = 18.41, p_bonf_ < 0.001, RTE_pre_ = 0.45, RTE_post_ = 0.55*), while performance on the regular keyboard remained unchanged (*ATS(1) = 0.784, p_bonf_ = 0.75, RTE_pre_ = 0.49, RTE_post_ = 0.51*). Finally, we observed a three-way interaction, Keyboard x Time x Group (*ATS(1) = 12.82, p < 0.001, RTE_REGpre_S_ = 0.75, RTE_MIRpre_S_ = 0.27, RTE_REGpost_S_ = 0.74, RTE_MIRpost_S_ = 0.35, RTE_REGpre_W_ = 0.7, RTE_MIRpre_W_ = 0.23, , RTE_REGpost_W_ = 0.71, RTE_MIRpost_W_ = 0.25*). Post-hoc test showed that typing performance on the mirrored keyboard improved after sleep (*ATS(1) = 25.27, p_bonf_ < 0.001, RTE_pre_ = 0.42, RTE_post_ = 0.58*) but not after wakefulness (*ATS(1) = 1.63, p_bonf_ = 0.81, RTE_pre_ = 0.48, RTE_post_ = 0.52*) thus confirming our hypothesis that sleep benefits motor adaptation consolidation. There was no change in the typing performance on the regular keyboard after retention neither in the sleep group (*ATS(1) = 1.26, p_bonf_ = 0.26, RTE_pre_ = 0.51, RTE_post_ = 0.49*) nor in the wake group (*ATS(1) = 3.65, p_bonf_ = 0.22, RTE_pre_ = 0.48, RTE_post_ = 0.52*). B) Sleep’s positive influence on motor adaptation did not interfere with the original task of regular typing. B) Typing performance on the regular keyboard in the different lists (List 1-3) before and after retention. Note that we used only words in List 1 for the training on the mirrored keyboard before retention; words in List 2 and List 3 were typed only on the regular keyboard before retention. A 2X2X2 mixed non-parametric test with two within-subject factors; *Lists* (List1, 2, and 3) and *Time* (before vs. after retention), and one between-subjects factor; *Group* (Sleep vs. Wake) showed only a main effect of *TIME* (*ATS(1) = 24.612, p < 0.001, RTEpre = 0.51, RTE_post_ = 0.49*) as performance increased after retention. There were no significant differences between the Lists (*ATS(1.63) = 0.267, p = 0.719, RTE_List1_ = 0.51, RTE_List2_ = 0.5, RTE_List3_ = 0.49*), or *Group* (*ATS(1) = 0.064, p = 0.801, RTE_Sleep_ = 0.49, RTE_wake_ = 0.51*). There was a significant interaction *Group X List* (*ATS(1) = 0.064, p = 0.801, RTE_Sleep_ = 0.49, RTE_wake_ = 0.51*). There was no significant interaction *Time X Lists X Group* (*ATS(1.99) = 0.495, p = 0.609*). C) No interference effects of sleep on regular typing. We compared the performance between sleep and the wake group on the words of List 1 (which participants trained on) before the retention on a word-by-word level. There was no significant difference in the performance between the groups (P>0.05). D) Comparisons between the different word lists (Lists 1, 2 and 3) after retention. Note that List 1 words were presented in MIR4, while MIR5 session contained words from lists 2 and 3. Results showed no effect of Group (*ATS(1) = 3.725, p = 0.122, RTE_sleep_ = 0.54, RTE_wake_ = 0.46*), but a main effect of *List* (*ATS(1.943) = 51.237, p < 0.001, RTE_List1_ = 0.74, RTE_List2_ = 0.58, RTE_List3_ = 0.18*). Performance was significantly higher in List 1 than List 2 (*ATS(1) = 62.719, p_bonf_ < 0.001, RTE_List1_ = 0.62, RTE_List2_ = 0.38*) and List 3 (*ATS(1) = 23.537, p < 0.001,* *RTE_List1_ = 0.74, RTE_List3_ = 0.26*), and List 2 than List 3 (*ATS(1) = 196.535, p < 0.001, RTE_List2_ = 0.74, RTE_List3_ = 0.26*). Finally, there was a significant interaction *Group X List* (*ATS(1.943) = 80.535, p = 0.031, RTE_List1_sleep_ = 0.82, RTE_List2_sleep_ = 06, RTE_List3_sleep_ = 0.2, RTE_List1_wake_ = 0.66, RTE_List2_wake_ = 0.56, RTE_List3_wake_ = 0.17*). Post-hoc Wilcoxon rank sum tests showed that for List 1 the performance was marginally higher for the sleep group as compared to the wake group (*W = 212, p_bonf_ = 0.081, 95% confidence interval [0.006 0.13]).* Each point in panel A represent the mean over one subject and the crossbars represent the mean over all subjects. REG1: Regular typing before retention, REG2: Regular typing after retention. MIR3: Mirrored typing before retention, MIR4: Mirrored typing after retention.

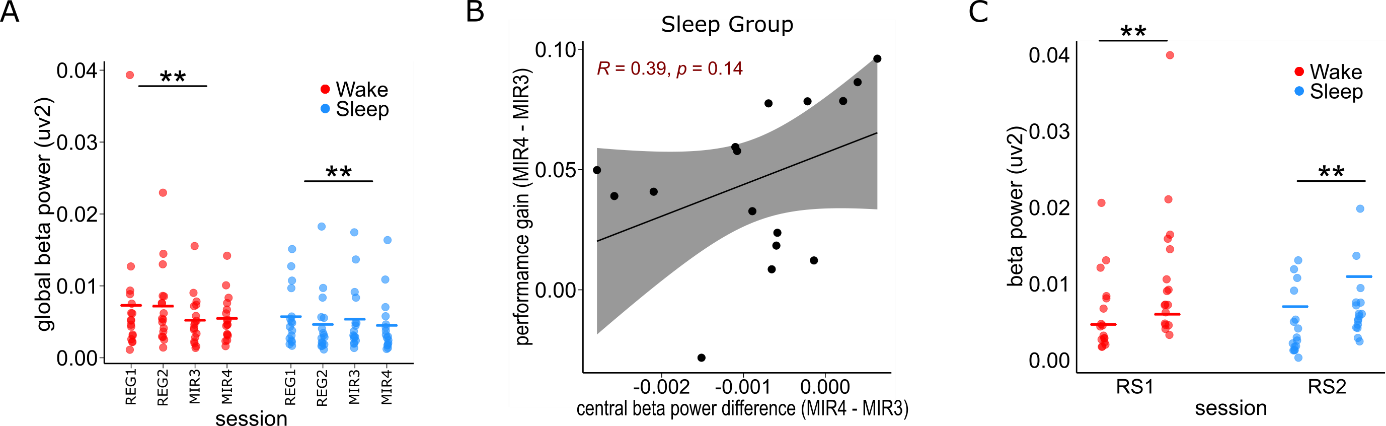

**Figure S3. Spatial and temporal aspects of movement-related beta band activity.** A) Global beta power differences between regular and mirrored typing. A 2x2x2 mixed non-parametric test with the two within-subjects Keyboard (REG vs. MIR) and Time (pre-retention vs. post-retention) and a between-subject factor Group (Sleep vs. Wake) showed that beta power was lower during mirrored typing (*ATS(1) = 13.57, p < 0.001, RTE_REG_ = 0.52, RTE_MIR_ = 0.48*). We found no effect of Time (*ATS(1) = 1.39, p = 0.23, RTE_pre_ = 0.51, RTE_post_ = 0.49) or Group (ATS(1) = 1.24, p = 0.27, RTE_Wake_ = 0.55, RTE_Sleep_ = 0.44*). However, there was a significant interaction Group x Time (*ATS(1) = 20.86, p < 0.001, RTE_SLpre_ = 0.49, RTE_SLpost_ = 0.4, RTE_Wpre_ = 0.53, RTE_Wpost_ = 0.58*), as beta power decreased after sleep (*ATS(1) = 20.29, p < 0.001, RTE_SLpre_ = 0.49, RTE_SLpost_ = 0.4*) but increased after wakefulness ( *ATS(1)= 7.815, p = 0.01, RTE_SLpre_ = 0.49, RTE_SLpost_ = 0.4*). There was no interaction *Keyboard x Time* (*ATS(1) = 0.77, p = 0.38*). We also observed a significant interaction Keyboard x Time x Group *(ATS(1) = 4.33, p = 0.04, RTE_REGpre_S_ = 0.51, RTE_MIRpre_S_ = 0.46, RTE_REGpost_S_ = 0.4, RTE_MIRpost_S_ = 0.4, RTE_REGpre_W_ = 0.55, RTE_MIRpre_W_ = 0.5, , RTE_REGpost_W_ = 0.62, RTE_MIRpost_W_ = 0.54*). Post-hoc tests showed that beta power decreased after sleep during both regular (*ATS(1) = 18.65, p < 0.001, RTE_pre_ = 0.56, RTE_post_ = 0.44*) and mirrored typing (*ATS(1) = 9.04, p = 0.01, RTE_pre_ = 0.54, RTE_post_ = 0.46*) but did not significantly differ from pre to post-wakefulness (Regular: *ATS(1) = 5.639, p = 0.07, RTE_pre_ = 0.47, RTE_post_ = 0.53 ; Mirrored: ATS(1) = 2.069, p = 0.6, RTE_pre_ = 0.48, RTE_post_ = 0.52*). B) The correlation between the change in beta power after sleep with the motor adaptation performance change from pre- to post-sleep. The change in beta power from pre-sleep to post-sleep mirrored typing session did not correlate with the change in behavioural typing performance on the mirrored keyboard. C) The effects of sleep on beta power are task related. A 2X2 mixed non-parametric test with Time as within subject variable and Group as between subject variable Comparing beta power during the eyes-closed resting state period before training (RS1) and before testing (RS2) revealed no difference in beta power between groups (*ATS(1) = 1.51, p = 0.22, RTE_Wake_ = 0.55, RTE_Sleep_ = 0.45*). However, there was a time effect as beta power increased from pre-retention to post-retention in both groups in the opposite direction of the task-related beta changes after sleep (*ATS(1) = 24.82, p < 0.001, RTE_SLpre_ = 0.35, RTE_SLpost_ = 0.54, RTE_Wpre_ = 0.42, RTE_Wpost_ = 0.68*). There was no significant interaction *Time X Group* (*ATS(1) = 0.52, p = 0.47*)

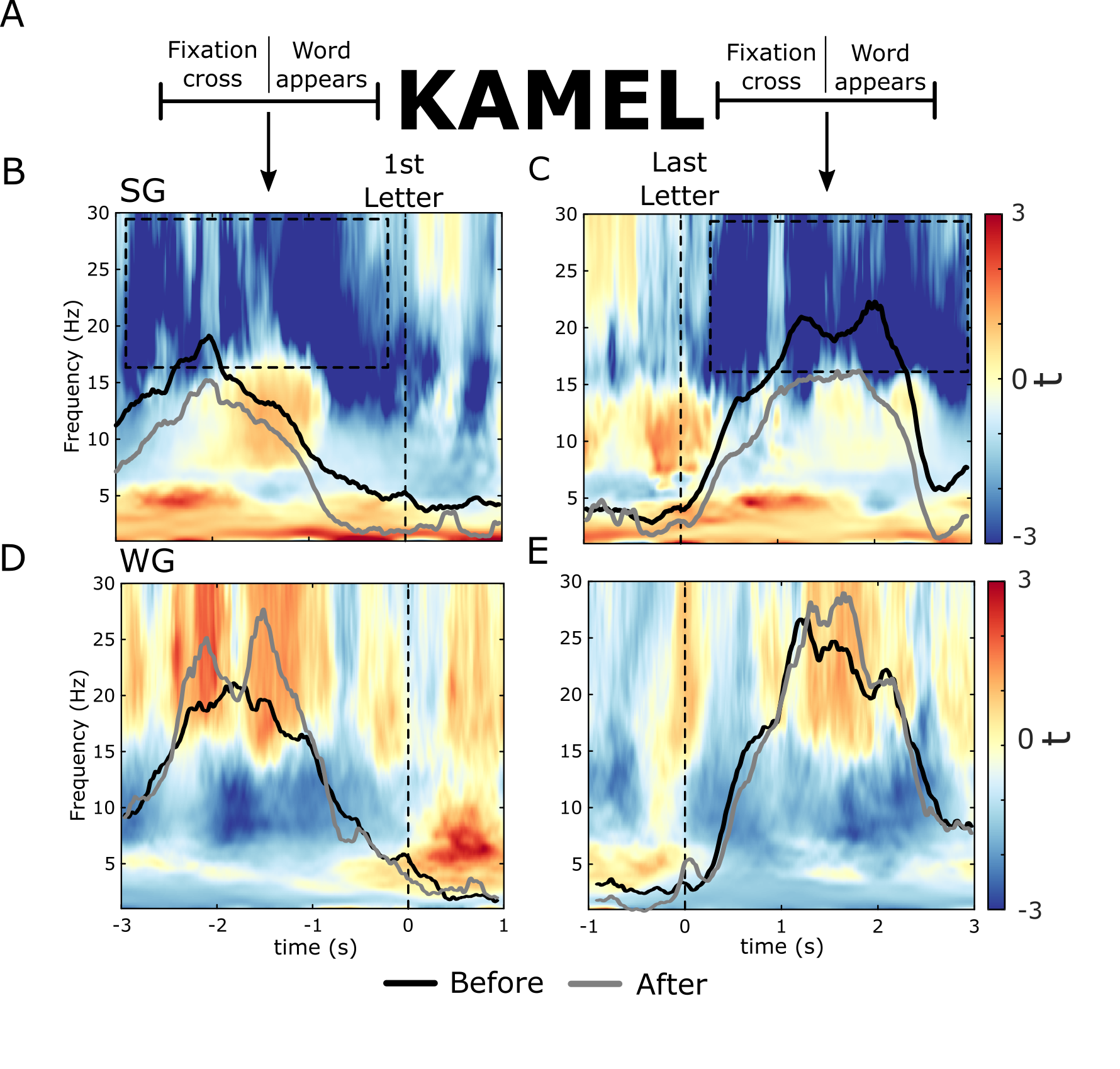
**Figure S3. Temporal dynamics of beta power during mirrored typing trials.** A) As participants had to type five letter words, to investigate pre-movement beta desynchronization and post-movement beta rebound periods, we looked at three second windows before typing the first letter of every word, and the three seconds following typing the last letter of each word, respectively. Note that in both cases, the analysed time window included the display of the fixation cross (1.5s) either before or after typing (for pre-movement and post-movement, respectively), and the appearance of a word on the screen, but did not include any typing activity. To confirm that there was no typing activity in this period, we measured the typing latency per session starting from the display of the fixation cross until typing the first letter of the word (desynchronization time) as well as the average time after typing the last letter of the word until typing the first letter of the following word (rebound window). The mean desynchronization time was 3.06s ± 0.48s (mean ± standard deviation) during MIR3 and 2.86s ± 0.69 during MIR4, while the mean rebound time was 4.17s ± 0.33 during MIR3 and 3.81s ± 0.38 during MIR4. B) Time-Frequency representations demonstrated a general decrease in the power of beta band following sleep during the pre-movement beta desynchronization periods (∑t(15) = -14006.95, p = 0.002) as well as (C) during post-movement beta rebound periods (t(15) = -18395.25, p < 0.001). We did not observe this decrease in beta power following a period of wakefulness (D-E). Note the black and grey lines on top of the time-frequency plots that indicate beta power averaged over C3 and C4, showing less beta desynchronization after sleep but not after wakefulness and lower rebound after sleep but not after wakefulness. The difference, however, was not statistically significant. REG1: Regular typing before retention, REG2: Regular typing after retention. MIR3: Mirrored typing before retention, MIR4: Mirrored typing after retention. SG: Sleep group, WG: Wake group.

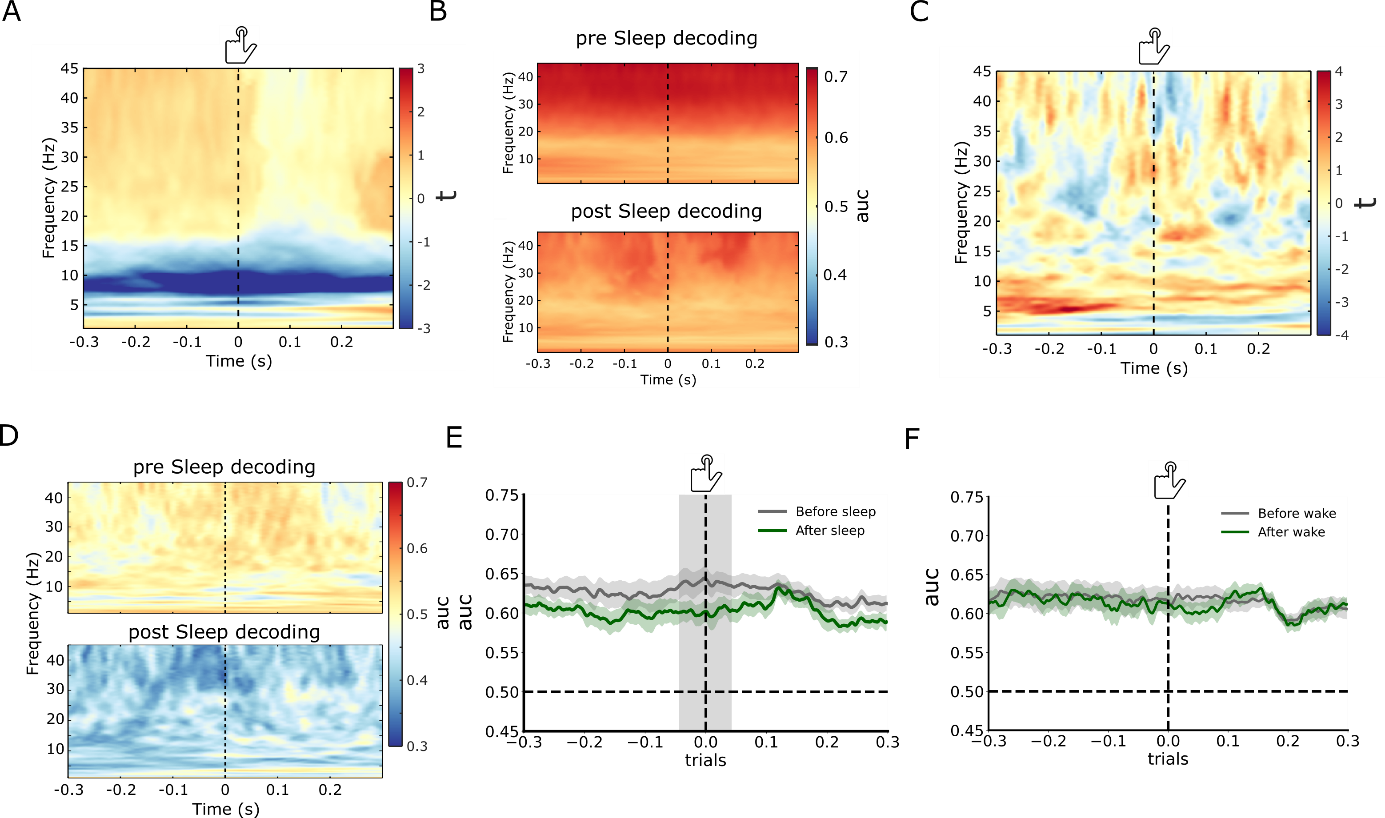

**Figure S5.** **Decoding results between regular vs. mirrored typing and correct vs. incorrect mirrored typing trials.** A) Frequency decoding on central electrodes in the wake group showed no change in beta band decoding accuracy (area under curve; auc) after retention. There was a decrease in decoding performance after wakefulness in the theta/alpha range that was, however, insignificant. B) Frequency decoding on central electrodes before and after sleep. Time-frequency decoding accuracy before and after sleep to show that beta band demonstrated the highest decoding performance before and after sleep. Note the decrease in decoding performance in the beta range after sleep. C) Frequency decoding of correct vs. incorrect mirrored typing trials in the wake group did not change from before to after retention (Post-wake – Pre-wake). D) A time-frequency representation of the frequency decoding on correct vs incorrect trials before and after sleep. E-F) According to our registered analysis plan, we decoded regular vs. mirrored typing using time-locked data averaged over all electrodes in the -0.3s to 0.3s window. E) Decoding performance dropped significantly after sleep (grey shading) in the time window *(-0.04s – 0.02s; ∑t(15) = -112.776, p = 0.012)*, but not after wakefulness (F).

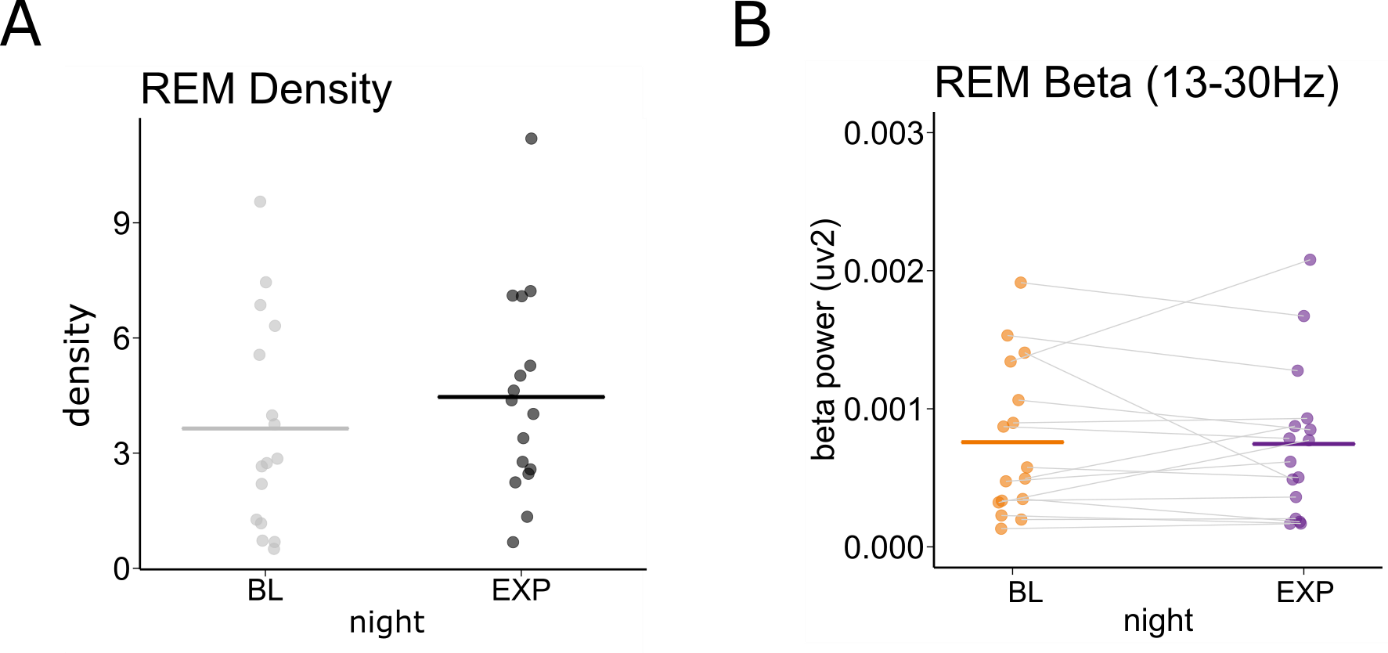

**Figure S6. REM beta power did not differ between baseline and experimental nights.** According to our registered analysis, we compared REM activity during the baseline night and the experimental night. A) No significant differences in REM sleep density (time spend in REM / Total sleep time) between the baseline night and the experimental night (ATS(1) = 1.36, p = 0.24, RTEBL = 0.55, RTE_Exp_ = 0.44). B) Beta power during REM sleep did not change from the baseline to the experimental night (ATS(1) = 0.03, p = 0.87, RTE_BL_ = 0.5, RTE_Exp_ = 0.5).

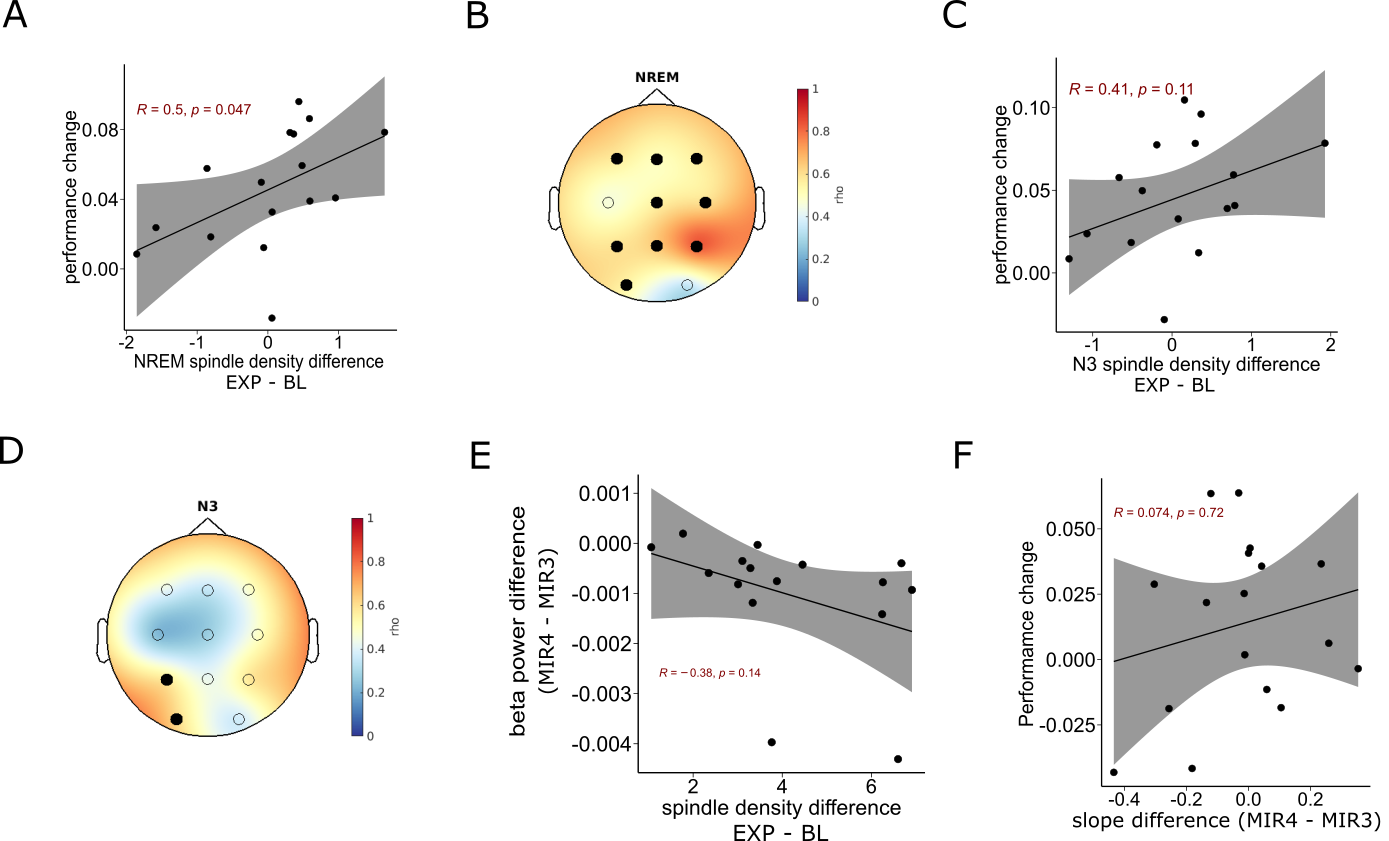

**Figure S7. Sleep results.** A-D) Fast spindle density (averaged over C3 and C4) correlation with the change in mirrored typing performance after sleep. A) When we measured fast sleep spindle density over the whole duration of NREM (N2 and N3) sleep we observed a significant positive correlation. B) The topography of correlation resembled that observed during N2 indicating a significant cluster of 9 electrodes (All except C3 and O2; *∑t(15) = 26.35, p = 0.003, d = 0.32*). C) N3 Fast sleep spindles density showed no significant correlation with the change in performance. D) Topography of electrodes exhibiting significant correlation of the change in fast spindle density with the change in mirrored typing performance (P1 and O1; *∑t(15) = 5.51, p = 0.03, d = 0.34*). E) The change in NREM stage 2 fast sleep spindle density from the baseline night to the experimental night did not correlate with the change in beta power from pre- to post-sleep during mirrored typing sessions. F) Spectral slope correlation with performance in the wake group. The change in slope from pre- to post-wakefulness did not correlate with the change in mirrored typing performance (MIR4 - MIR3). REM: rapid eye movement sleep. R: Pearson’s correlation coefficient. Rho: spearman’s correlation. BL: baseline night, EXP: experimental night.
